# Decision-making in a synthetic cell: the limits of biological computation

**DOI:** 10.1101/2020.05.29.116467

**Authors:** Ferdinand Greiss, Shirley S. Daube, Vincent Noireaux, Roy Bar-Ziv

## Abstract

We measured the dynamics of decision-making by a minimal bistable gene network integrated in a synthetic cell model, free of external perturbations. Reducing the number of gene copies from 10^5^ to about 10 per cell revealed a transition from deterministic and slow computation to a fuzzy and rapid regime dominated by single-protein fluctuations. Fuzzy computation appeared at DNA and protein concentrations 100-fold lower than necessary in equilibrium, suggesting rate enhancement by co-expressional localization. Whereas the high-copy regime was characterized by a sharp transition, hysteresis and robust memory, the low-copy limit showed incipient strong fluctuations, switching between states, and a signature of cellular individuality across the decision-making point. Our work establishes synthetic cells operating rapidly at the single molecule level to integrate gene regulatory networks with metabolic pathways for sustained survival with low energetic cost.

**One Sentence Summary:** Decision-making in a synthetic cell can be slow and precise or rapid and probabilistic by reducing the number of computing molecules by five decades down to single-molecule fluctuations.

## Main Text

Making well-informed decisions takes time and energy. The fundamental connection between those three aspects is exemplified by recognition memory in cognitive tasks (1), optimality in molecular recognition as an error-correction mechanism (2,3), and cellular adaptation in chemo-sensing (4). Decisions on a cellular level are made by regulatory proteins that integrate information from the environment and elicit a response by modulating RNA and protein production. In each cell, the copy number of regulatory molecules could vary between one to a few hundred, subjecting the information processing to random fluctuations (5,6). The fluctuations can stem from production and degradation of proteins, and the binding and unbinding of proteins to and from their regulatory sites on DNA. Theoretical considerations suggest that a decision becomes exponentially more stable as the copy number of regulatory elements, hence metabolic load, increases (7,8).

Decision-making of genetic regulatory networks (GRN) should be precise and deterministic when the process is averaged over many computing molecules. However, in the small-number limit, fluctuations in gene-expression with a few computing molecules, hence little averaging, may reduce the precision and lead to fuzzy computation (Fig. 1A-B). By measuring the characteristics of a GRN in the high and low copy number regime, we addressed the following questions: Can we measure the transition from deterministic to fuzzy decision-making dynamics driven by a GRN? Can we recognize a fundamental tradeoff between the two regimes? And how do single-molecule fluctuations influence the decision-making?

**Fig. 1.**
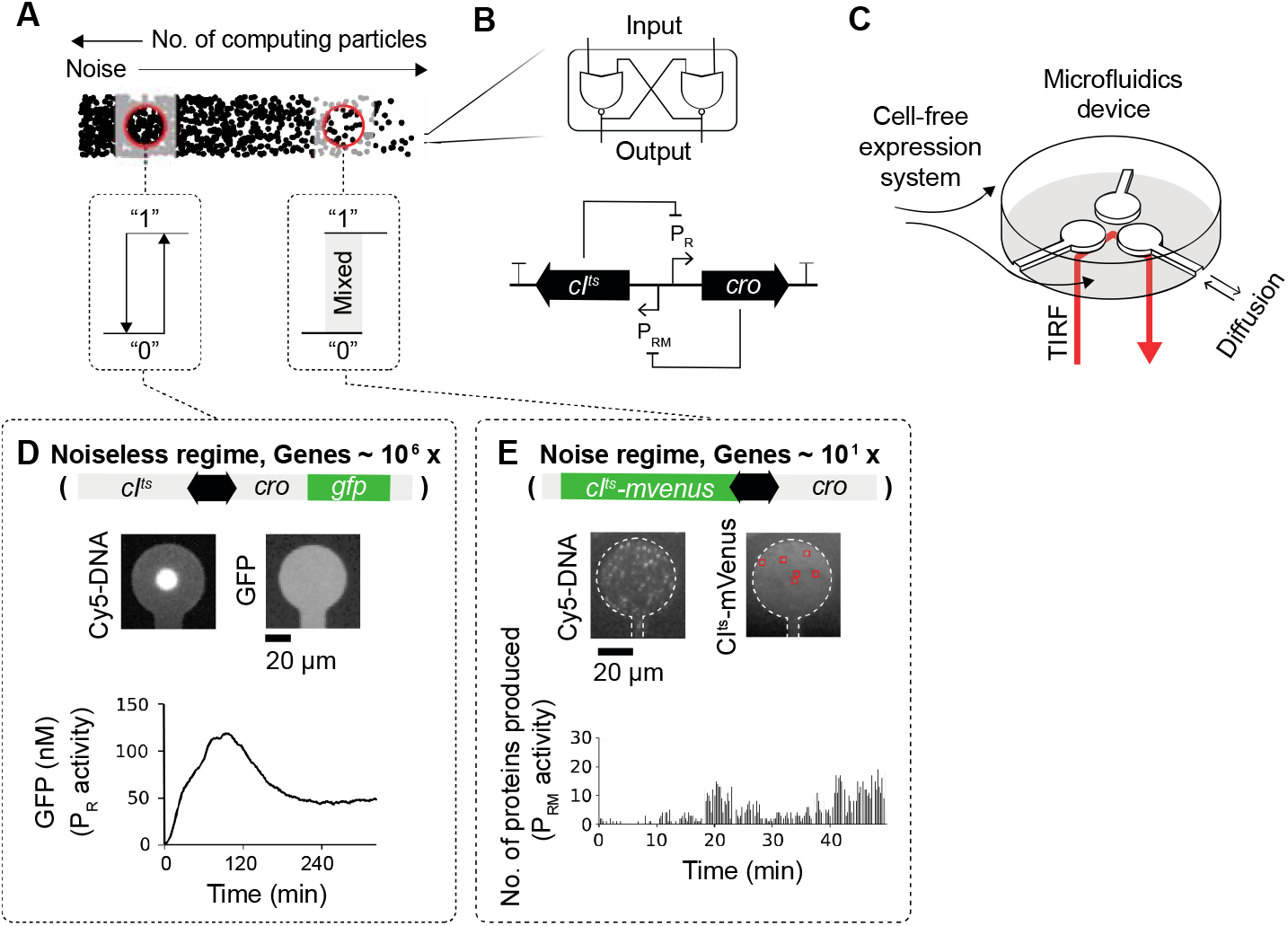
Biological computation in a synthetic cell model at high and low gene numbers. (**A**) Number of computing particles influence decision-making by an effective noise level: a bistable GRN responds to an input with a binary output in the noiseless regime at large numbers, or by a fuzzy-logic computation with mixed outputs in the noise regime at small numbers. (**B**) Scheme of the minimal bistable GRN based on CI and Cro transcription factors repressing each other’s promoter with analogy to an electronic latch circuit. (**C**) Scheme of a microfluidic device with three compartments as synthetic cell models. (**D**) Cy5-DNA assembled in a compartment at high density gives rise to GFP expression reporting on the P_R_ activity. **(E)** Cy5-DNA at single molecule resolution inside the compartment. Single CI^ts^-mVenus were integrated over the compartment to give number of proteins produced in 15 seconds, reporting on the overall P_RM_ activity.

In order to address these questions experimentally, a programmable cell-free system of gene-expression reactions with constant protein turnover is required (9). To that end, we built synthetic cell models devoid of extrinsic noise stemming from DNA replication, genetic cross-talk, or cell size variation (10–12). Each cell had a DNA compartment with a 20 μm radius and 3.5 μm height, connected by a thin capillary (L=90 μm and W=7 μm) to a reservoir of cell-free extract to support transcriptiontranslation (TXTL) reactions (Fig. 1C, Fig. S1) (13). The compartment volume was 3.8 10^3^ μm^3^, roughly 1000-fold larger than a typical bacterial cell. The capillary allowed free diffusion in and out of the compartment, creating TXTL dynamics with an apparent protein lifetime of *τ = πR^2^L/WD* = 8 min (13), with D = 30 μm^2^/s as the typical protein diffusion coefficient. We used an elastomer to create the compartments, bonded to a coverslip to detect the synthesis of fluorescent reporter proteins from the DNA attached on the glass substrate (Methods).

We based our genetic model system on a bistable GRN built from the core elements of the lambda bacteriophage regulatory network (Fig. 1B). In its native environment, this GRN controls the phage’s decision to lyse the cell or integrate its genome into the bacterial host genome thus entering the lysogenic state. This state can be maintained for more than 10^5^ generations by the presence of only a few hundred copies of proteins before switching back to the lytic phase (10,14–17). The minimal GRN consists of two transcription factors, CI and Cro repressors, which mutually inhibit each other’s production by binding to their respective promoters (P_R_ and P_RM_) (Fig. 1B) (16). By analogy to digital electronics, an ideal bistable GRN can be viewed as a latch circuit that activates either of two promoters and remembers the active promoter until toggled (Fig. 1B). CI has been shown to be the main regulator, responsible for entering and maintaining the lysogenic state, while Cro buildup serves a tipping point to decisively and irreversibly enter the active P_R_ promoter (lytic) state (18). This inherent asymmetry between CI and Cro is due to the promoters’ architecture including auto-inhibition and auto-activation loops (16). To toggle the promoter activities, we used a temperature-sensitive CI (CI^ts^) mutant that tunes its deactivation rate with a rise in temperature from 30 °C (no deactivation) to 41 °C (fast deactivation) (19,20).

We immobilized the DNA constructs with the bistable GRN on the surface of the compartments. At the high-density regime, we packed roughly 10^5^-10^6^ copies (21) in a DNA brush patterned on a circle of 14 μm diameter (Methods). We placed the gene coding for green fluorescent protein (GFP) in tandem with the *cro* gene, as a reporter of the activity of the strong P_R_ promoter (16,20) (Fig. 1D), and recorded the signal using epi-fluorescence microscopy. We observed a smooth and noiseless GFP signal, peaking after ~2 hours and slowly reaching steady-state values after two more hours (Fig. 1D).

For the low-copy regime, we immobilized an average of ~20 copies of the bistable GRN in each compartment, which amounted to a very low effective DNA concentration of ~10 pM (Fig. 1E and Fig. S2). To allow the detection of single proteins, we fused the *cI^ts^* gene under the weak P_RM_ promoter (16,20) to the fast-maturing fluorescent protein mVenus (22), and circularized the DNA constructs to minimize degradation by exonucleases in the *E. coli* extract (Methods, Fig. S3, Table S1). We employed total internal reflection fluorescence (TIRF) microscopy with a penetration depth comparable to the typical dimension of DNA molecules to observe localization of the proteins close to the coverslip surface.

After introducing the cell-free extract, we could observe the appearance of discrete CI^ts^-mVenus fluorescent spots that were single-peaked and rapidly bleached with a lifetime of ~4 sec (Fig. S4). The fast bleaching removed all newly produced proteins that localized to the surface. We could not reliably detect repeated binding of proteins to specific locations due to nonspecific adsorption on the surface upon heatinactivation of CI^ts^-mVenus (Fig. S5). Therefore, we integrated the number of produced proteins inside the compartment independently of the location into 15 sec time intervals, reporting on the overall P_RM_ activity of the minimal cell model (Fig. 1E) (20,23,24). In sharp contrast to the high-density regime, we observed strong fluctuations of protein production rates that reached a first maximum after 20 min.

Because we optimized the cell extract such that the basal protein production rate varied by no more than ~10% for 31-41 °C (Fig. S6), any change in signal properties as a function of temperature could be attributed to a response of the GRN to a change in CI^ts^ deactivation rate. Based on the GRN architecture studied in bacteria (10,16,20), we anticipated initial low P_RM_ and low P_R_ activities at low temperature, where stable CI^ts^ first represses P_R_ and then self-represses P_RM_ at the mutual promoter sites (Fig. 2A, state 1). An increase in CI^ts^ deactivation rate at higher temperature should reduce the occupancy of CI^ts^ at the promoters, leading to selfactivation of CI^ts^ and gradual production of Cro from P_R_ (Fig. 2A, state 2). At higher temperatures, complete deactivation of CI^ts^ leads to fully active P_R_ (Fig. 2A, state 3).

**Fig. 2.**
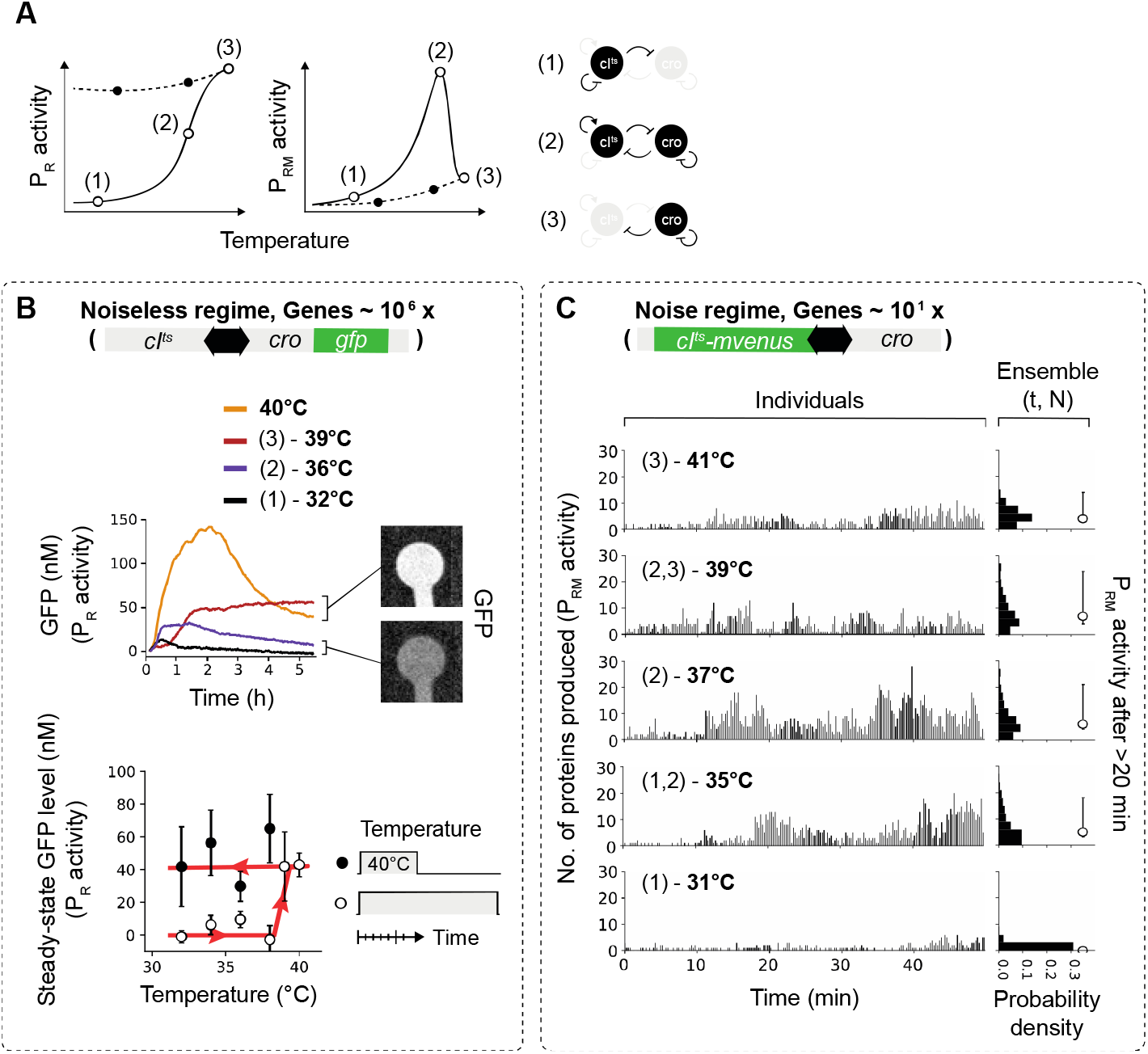
Bistable GRN and decision-making at high and low gene densities. (**A**) Illustration for the expected output of P_R_ and P_RM_ activity as a function of temperature, based on the known molecular feedbacks of the GRN. **(B)** Dynamics of GFP expression reporting on the P_R_ activity at high DNA density (upper panel). Solid lines depict average dynamics, N=24. (1), (2), (3) relate to molecular states of the bistable GRN at each temperature, as in (A). Steady-state GFP levels averaged over individual compartments (lower panel). Temperature was either kept constant throughout the measurement (open circle) or after a 40 °C initial incubation step (closed circles). Error bars show standard deviation (SD) of compartments. (**C**) Number of CI^ts^-mVenus produced in 15 seconds to give production rates (left column) and ensemble distribution over time t and compartments N after >20 min (right column) in the low-copy regime. Circle with error bars give the median and 32th to 68th percentile of the ensemble production rates.

At the high-density regime with GFP, hence Cro, reporting on P_R_ activity, no signal was recorded at 31 °C, indicating an active P_RM_ promoter (Fig. 2B, upper panel, Fig. S7, and Fig. 2A, state 1). We observed a decisive transition between 38 °C and 39 °C to high GFP steady-state levels, switching to active P_R_ promoter (Fig. 2A, state 3). To check the stability of the activated P_R_ promoter, we first expressed the proteins in the monostable region at 40 °C for 1 h, followed by a drop to lower temperatures. P_R_ activity was reliably maintained (Fig. 2B, lower panel and Fig. S7), independently of the compartment geometry and hence protein lifetime (Fig. S9 and Fig. S8). Control experiments with a GRN lacking the *cro* gene (a monostable GRN) resulted in a similar initial response to temperature variations, but with overall lower GFP levels due to the lack of commitment to the P_R_ promoter by Cro, as was already observed in bacteria (Fig. S9) (18), suggesting that Cro was required to obtain decisive switching (Fig. 2B, lower panel). Overall we observed a deterministic response to the temperature input, most likely due to low noise levels, with precise but slow dynamics on a characteristic time-scale of ~1 h (Fig. S10), consistent with the bistable feature of the GRN as an ideal memory device. Solution experiments reconstituted a similar temperature response of the different GRNs, but could not identify a decisive switching temperature since the closed system accumulated proteins without turnover, and hence did not reach steady-state dynamics (Fig. S11).

In the low-copy regime, with CI^ts^-mVenus reporting on the P_RM_ activity, the overall production rates of an ensemble of compartments (Fig. S12, Fig. 2C, right column and Table S2) increased (Fig. 2A, state 2) and decreased again at higher temperature as anticipated (Fig. 2C and Fig. 2A, state 3). We observed variability between compartments only in the 35-39 °C range as computed from the standard deviation of time-averaged production rates (Fig. S13). In this range, production rates on the time scale of many minutes (~5 min) waned and grew again (Fig. 2C, Fig. S10, Fig. S12). In the absence of Cro, we expected to observe a P_RM_ activity without inhibition by Cro, that is no display of spontaneous transitions between the two promoters. Supportively, the temperature response for the monostable GRN was similar to the bistable GRN (Fig. 3A,B and 2C, respectively and Fig. S12), but occurred across a narrower temperature range with higher median production rate at 37 °C (Fig. 2C, Fig. 3A, and Table S2), and reduced variability at 37 °C and 39 °C (Fig. S13). A control experiment of a monostable GRN with the wild-type *cl* gene displayed almost no temperature response (Fig. S14 and Table S2). To exclude the possibility that the high concentration of the TXTL machinery in the cell-free extract (25) governed the GRN dynamics, we replaced the P_RM_ by a consensus promoter sequence and added a ribosomal binding site (RBS) to the natural leaderless mRNA (Fig. S15) (26). We observed a strong increase in production rates in both cases. These control experiments supported P_RM_ activity as the rate limiting step of the GRN, also validating CI as the main regulator of P_RM_. We also found no dependence of protein production rates on the slightly varying number of DNA molecules in the low-density compartments (Fig. S2). The data therefore suggests that in the bistable GRN either one of the two promoters was randomly activated and could spontaneously transition to the other promoter, but only at the 35-39 °C temperature range (20). We also note that the variability occurred in a broad temperature range close to the switching point of the high-density regime.

**Fig. 3.**
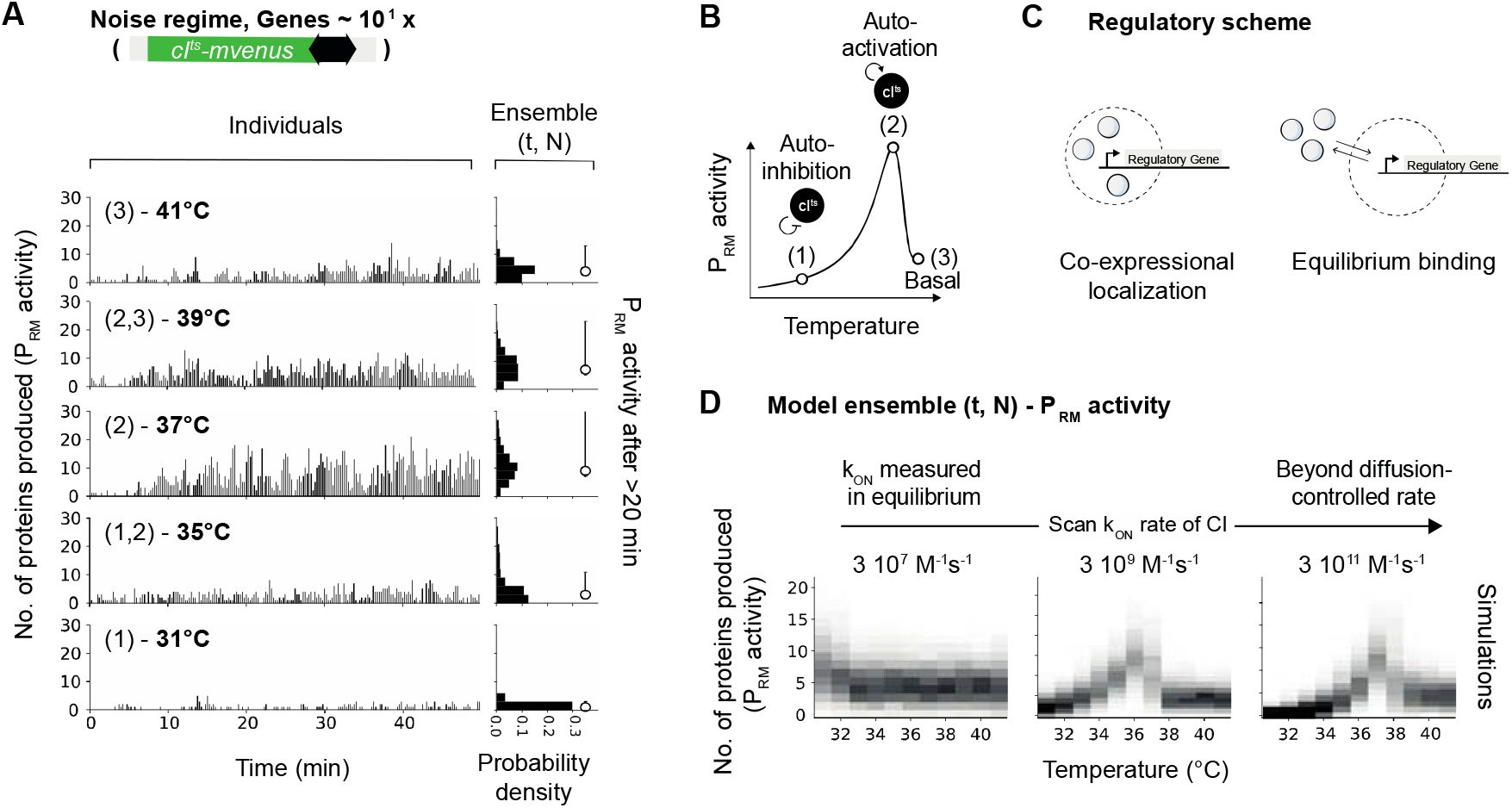
Monostable GRN and co-expressional localization at the low-density limit. **(A)** Production rate of CI^ts^-mVenus proteins in 15 seconds inside individual compartments (left column) and ensemble production rates (right column) as in Fig. 2C. Circle with error bars give the median and the 32th to 68th percentile of the ensemble production rates. (**B**) Illustration of the expected P_RM_ activity with temperature in the monostable GRN and the corresponding states. (**C**) A protein repressor synthesized in proximity to its DNA binding site, reach higher effective concentrations by coexpression localization (left scheme) than in an equilibrium scenario due to diffusion away from the DNA (right scheme). **(D)** Ensemble production rates obtained from stochastic simulations with a scan in the CI^ts^ binding rate to the DNA binding sites.

Whereas computational speed seemed to be traded for precision at an energetic cost of ~10^6^ proteins in the high-density regime, fuzzy computation led to increase in speed with only a handful of proteins in the low-density regime (Fig. S10). Still, we wondered how decisions could be realized in the low-density regime considering the low DNA and protein concentration. We estimated the total number of proteins produced in 50 min to be ~1000, which amounted to an upper-limit concentration of ~50 pM. With DNA at 10 pM, the concentrations were below the measured CI binding affinities of ~3 nM (27) and ~50 nM (12), *in vitro* and *in vivo*, respectively, suggesting that equilibrium binding considerations could not account for the binding of nascent CI^ts^ to its DNA binding site. Considering the typical binding rate of CI to its operator sites of k_on_ ≈ 3 10^7^ M^-1^s^-1^ (28), it would take a single protein much more than 20 min to bind and modulate production rates, slower than the observed time to reach steady-states. These considerations could be reconciled by a scenario in which regulatory proteins are localized to the DNA during production and kept close to their operator sites with an increase in local effective concentration (Fig. 3C). Since our system did not allow us to observe promoter activity, mRNA production, and protein synthesis at the same time and space, we corroborated this notion by stochastic simulations and found that they reproduced the experimental results only at values 100-fold higher than previously measured CI k_on_ rates in equilibrium (compare Fig. 3A to Fig. 3D and Table S3). Likewise, CI^ts^ repressed the production of GFP in solution experiments more efficiently when produced on the same plasmid than on a separate plasmid (Fig. S16). This notion of spatial coupling between protein synthesis and DNA-binding as an enhancement mechanism for site location was also discussed in a bacterial context (29) but was not demonstrated *in vitro* so far.

Next, we sought to find the source for the fuzzy decision-making in the low-density regime. The Fano factor was computed as a measure of fluctuations in CI^ts^-mVenus production rates in single compartments after 20 min (the variance/mean = 1 for a Poissonian process) and plotted in Fig. 4A. The Fano factor was close to 1 and constant with temperature for the wild-type monostable GRN. For both mono- and bistable GRNs, the Fano factor was greater than 1 around 37 °C, and correlated with the time-averaged production rates (Fig. 4A, upper row, Fig. S17). In addition, the Fano factor of the bistable GRN had a larger spread between 35-39 °C in the range of large variability. The data indicated that the regulation of the P_RM_ activity by CI^ts^-mVenus strongly fluctuated only around 37 °C. Here, the fast inactivation and therefore lower occupancy of CI^ts^-mVenus at the P_R_ promoter seemed to reduce the stability of the auto-activation of P_RM_ activity by CI^ts^-mVenus (Fig. 4B and Fig. S18); hence a lower occupancy at the P_R_ promoter allows the leaky production of Cro to induce the observed variability and spontaneous transitions between promoters in the bistable GRN, resulting in fuzzy decision-making.

**Fig. 4.**
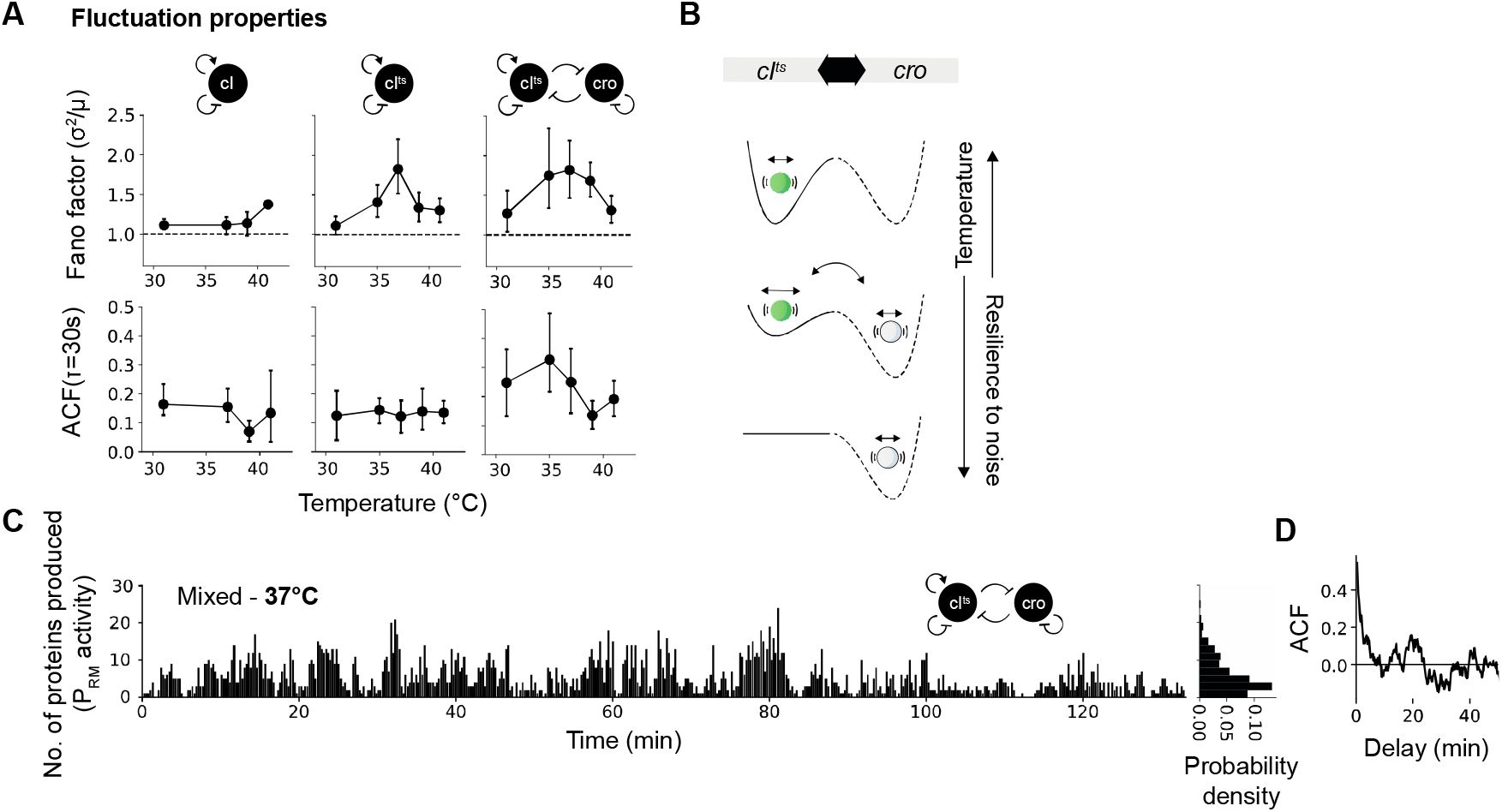
Fluctuations and spontaneous transitions at the low-density limit. (**A**) Fano factor (variance over mean) with Poissonian process (dashed line) for the GRNs at various temperatures (upper row). Amplitude of temporal auto-correlation function (ACF) of fluctuating production rates (Fig 2C, 3A, Fig S16) in individual compartments at τ = 30 s. Error bars were bootstrapped and show SD of compartments. (**B**) Scheme of noise, active promoter transitions and memory in the bistable GRN. Short-term fluctuations in gene expression (circle within a well) originate from local noise. Long-term fluctuations can originate from spontaneous promoter transitions with the deactivation of CI^ts^. (**C**) Number of proteins produced in 15 seconds for long-term experiment of the bistable GRN at 37 °C. (**D**) The ACF of the trace in (C).

To extract typical correlation time scales in the fluctuating production rates of CI^ts^-mVenus after 20 min, we computed the autocorrelation functions (ACF) in each compartment. The ensemble-averaged ACFs decayed within a few minutes as expected for the random protein synthesis from ~20 DNA molecules (Fig. S17) (23). However, the ACF amplitude at a delay of τ = 30 seconds had a higher amplitude for the bistable GRN than the wild-type and monostable GRN at 39 °C and below (Fig. 4A, lower row and Fig. S17). This may indicate more correlated fluctuations of CI^ts^-mVenus production rates for the bistable GRN as was observed for a close bifurcation point in other complex dynamic systems, and can be explained by a reduced recovery rate from perturbations (30). Here, molecular perturbations may stem from the leaky production of Cro from P_R_ within the region of bistability (Fig. 4B). To get better statistics on the long-term fluctuations that could last for many minutes in the bistable GRN (Fig. 2C), we extended the duration of the experiment to 130 min at 37 °C (Fig. 4C). We observed highly fluctuating production rates with repeating signatures of protein bursts, each lasting for several minutes (Fig. 4D and Fig. S19). A strongly damped periodic signal could be observed in individual ACFs with a period of ~10 min, but averaged out when ensemble-averaged (Fig. S18).

We established a minimal decision-making GRN, with and without variability and spontaneous transitions, in a synthetic cell devoid of any extrinsic noise. Dynamics could be monitored at the low and high copy number limits of genetic computation, demonstrating a clear tradeoff between slow and precise versus fast and fuzzy. We discovered evidence for enhanced rates by co-expressional localization (Fig. 3) (29), a nonequilibrium mechanism that seems to be essential for realizing single-molecule regulation in our minimal cell models.

## Acknowledgments

We thank M. Levy and O. Vonshak for fruitful discussions and help with experiments; L Tunik for help in the fabrication facilities; N. Stern, M. Schwarz-Schilling, and T. Tlusty for helpful comments on the manuscript; and J. Götz for the brilliant support on all levels throughout the project. We thank D. Garenne for his participation in the preparation of the TXTL system used in this work. pGEX iLOV was a gift from John Christie (Addgene plasmid #26587; http://n2t.net/addgene:26587; RRID:Addgene_26587).

## Author contributions

F.G. designed the system, performed experiments and analyzed the data. All authors discussed data and wrote the manuscript.

## Supplementary Materials

Materials and Methods

Figures S1-S19

Tables S1-S3

