## Supplementary Materials for "Decision-making in a synthetic cell: the limits of biological computation"

### *Materials and Methods*

**Cell-free transcription-translation.**

Cell-free expression was carried out using an *E.coli* TXTL system (myTXTL, Arbor Biosciences) as previously described (25,31).

**Reagents.**

All primers were ordered from IDT. All PCRs were performed with the KAPA HiFi HotStart ReadyMix (Kapa Biosystems, Roche). All PCR products were purified with Promega Wizard SV-Gel and PCR Clean-Up. All plasmids were derived from the pBEST backbone (25) with Ampicillin selection marker and ColE1 origin of replication. Cloning was performed with a *E. coli* strain (DH5α) and plasmids were extracted with spin column purification (Promega).

**Assembly of gene GRNs.**

The GRNs with $cI^{ts}$, $O_R3$, $O_R2$, $O_R1$, and *cro* were directly isolated from lambda phage DNA (methylated lambda phage DNA, Sigma) using primers P3 and P4 for transfer-PCR into the pBEST vector to obtain a plasmid with the minimal bistable GRN and the RBS (UTR1) and GFP sequence in a bicistronic design with the *cro* gene (Table S1) (25). This plasmid served as the template for all further modifications that were introduced with PCR using phosphorylated primers, ligated (T4 Ligase, ThermoFisher Scientific), and directly transformed into bacteria. Constructs were sequenced by the sequencing service of the Weizmann Institute.

The $cI^{ts}$-*mvenus* construct was generated by fusing the C-terminus of the $cI^{ts}$ gene to the codon-optimized and truncated *dmvenus* sequence (25) with the point mutations for fast maturation and the monomeric state (L64V and A202K) resulting in *dmvenus-NB* (also known as SYFP2). Here for simplicity, we termed the fluorescent protein mVenus. The two proteins were genetically fused by a flexible linker (KRAPGTS, AAGCGAGCTCCCGGGACCAGC). The gene fragment of *mvenus* and linker was ordered as DNA fragment (gBlock, IDT) and after removing the *gfp* gene, cloned into the plasmids of the mono- and bistable GRNs using transfer-PCR.

**DNA protection.**

Linear DNA is quickly degraded in cell-free extract without protection from enzymatic activity. RecBCD is essential for recombination in bacteria, but is known as key factor in the degradation of linear and single-stranded DNA. Commonly, the DNA-mimicking lambda bacteriophage protein GamS is supplemented in high amounts (~1 μM) to outcompete RecBCD (32). But, to further improve the stability of DNA at the low-density regime, we devised a protocol to avoid using linear DNA, that is to prepare fluorescently labeled and biotinylated DNA without open ends at high concentrations for controlled surface immobilization.

The assembly protocol for the circular and hairpin DNA was the following:
Two PCR mixtures were prepared according to the manufacturer's protocol. The first mixture contained the template, one phosphorylated primer pP1, and one primer P2 (that is the sequence of pP2 without phosphorylation and overhang complementary to P1) with internal modification, e.g. biotinylated thymidine residue. The second PCR mixture contained the same template, a phosphorylated primer pP2 and a primer P1 (that is the sequence of pP1 without phosphorylation and overhang complementary to P2) with internal modification, e.g. internal Cy5 label. After PCR purification, the products hold two almost identical dsDNA fragments except for the end modifications (P1, P2). For the final step, the two dsDNA fragments were mixed (2.5 μg of each) and incubated with lambda exonuclease (5 units, NEB) in lambda exonuclease reaction buffer at 37 °C for 1 h to selectively degrade the end-phosphorylated (pP1, pP2) ssDNA parts (Fig. S3). The exonuclease was heat inactivated for 15 minutes at 75 °C. The solution was cleaned and stored at -20 °C.

**Closed system gene expression.**

Calibrations (Fig. S6) and GRN control experiments (Fig. S11) were done in a real-time PCR system (StepOnePlus, Applied Biosystems) that allowed simultaneous gene expression at 6 different temperatures. The system was calibrated with purified GFP in 1x phosphate-buffered saline (PBS) to linearize the readout. All solution experiments were conducted in a volume of 10 μl with 1 nM of plasmid. The conditions for cell-free gene expression were optimized to reduce temperature variations in basal GFP expression. For this batch of extract, a level of $Mg^{2+}$ and polyethylene glycol 8000 at 10 mM and 4 % respectively, gave ~10% expression differences in the range between 30 °C and 41 °C (Fig. S2).

For the transient temperature variations in the solution experiments (closed system), we further implemented iLOV as fluorescent reporter that was shown to be oxygen-independent, temperature insensitive, and quickly maturated (Fig. S11) (33). However, we found that the brightness was not high enough to also use it as reporter for gene expression in the open systems.

At constant temperatures, all solution experiments were conducted with a plate reader (ClarioStar Plus, BMG Labtech) with 1 nM plasmid in a volume of 10 µl (Fig. S3 and Fig. S16).

**Fabrication of microfluidic chips.**

The mold for the microfluidic chip was fabricated using standard clean-room equipment and SU-8 photoresist (MicroChem, Newton, MA). A clean 4” silicon wafer (4”, 0.525 mm thickness, <100>, p-type, University Wafers) was carefully covered with hexamethyldisilazane (HMDS, Transene Company) and incubated for 30 sec to promote adhesion of the SU-8 photoresist. After incubation, the wafer was spun (PWM32, Headway Research Inc., Garland, TX) for 30 seconds at 3000 rpm with a ramp of 1000 rpm/sec. The wafer was coated (compartment layer) with a 3 µm layer of SU-8 2002 (stage 1: 7.5 seconds at 500 rpm with acceleration of 100 rpm/sec, stage 2: 30 seconds at 750 rpm with acceleration of 300 rpm/sec). The photoresist was processed according to the manufacturer’s protocol with a heating plate and exposed after careful alignment using 5” chrome photomasks (Nanofilm) and mask aligner (Karl Suss Ma6/BA6, Garching, Germany). The patterned photoresist was treated according to the manufacturer’s protocol with a heating plate omitting the hard baking step. The second layer (feeding layer) with 65 µm of SU-8 3050 was spun on the wafer (stage 1: 7.5 seconds at 500 rpm with acceleration of 100 rpm/sec, stage 2: 30 seconds at 2000 rpm with acceleration of 300 rpm/sec) and again processed according to the manufacturer’s protocol with the final hard baking step (slow ramping from room temperature to 150 °C with slow cool-down to room temperature without removing the wafer from the heating plate).

The SU-8 mold with 9 microfluidic chips was covered with polydimethylsiloxane (PDMS, 10:1 ratio of polymer and curing agent, Sylgard 184, Dow Corning) in a petri dish, air bubbles were removed under vacuum for ~2 h, and baked overnight at 70 °C. The cured PDMS block was gently peeled off the wafer and cut in pieces. The holes for inlet and outlet were punched on a cutting mat (0.75 mm diameter, Welltech Labs), thoroughly cleaned with isopropanol and blow dried with nitrogen. Fresh 170 µm coverslips (No. 1.5H, Marienfeld) were assembled in a Teflon holder, boiled in 96% ethanol for 10 minutes and transferred to a clean beaker with 1:3 $NH_3$(25%):$H_2O$ to be heated to 70 °C. Once heated, one part of $H_2O_2$ was supplemented and boiled for another 10 min. The coverslips were transferred into clean $H_2O$ and blow dried with nitrogen for storage. Coverslips and PDMS parts were plasma treated at 35 watts for 30 seconds with 1.5 sccm $O_2$ inflow (Plasma System and GCM-200, March Plasmod) and brought in firm contact immediately after. The assembled chip was finalized by baking at 70 °C overnight.

**DNA patterning for low and high gene density.**

An amount of 0.3 mg of biocompatible photoactivatable polymer solution (termed "DAISY") (34) was dissolved in 1.5 mL acetonitrile (HPLC graded) and flushed onto four chips through PTFE tubing (Cole Parmer). The microfluidic chips were incubated for 20 minutes to form a self-assembled monolayer on the surface. The solution was then successively exchanged by flushing 1 ml of 100%, 50%, 25%, and 0% of acetonitrile (HPLC graded, Bio Lab LTD, Israel) and $H_20$ mixture. The unprotected DAISY chains were blocked prior to UV illumination using Methyl-PEG$_4$-NHS (ThermoFisher Scientific, 5 mg/ml in 250 mM borate buffer, pH 8.6) for high gene density. The chip was thoroughly washed with $H_2O$ after a 5-minute incubation.

Photolithography was then performed in a UV cube (UV KUB, Kloé, France) for low gene density or on a standard optical microscope (Axiovert 200M, Zeiss) with a mercury arc lamp (X-Cite Series 120Q, Excelitas Technologies Corp, USA) for high gene density and spatial patterning. For low gene density, the chip was exposed continuously for 100s at 100% power. For high gene density, the excitation spectrum was filtered using a 365±10 nm band-pass filter (Chroma). A 200 µm spherical aperture (P200D, Thorlabs) was introduced as field stop into the microscope and a second adjustable spherical aperture was used to reduce the numerical aperture, hence reducing the influence of defocusing. The pattern was then passed through an 40x/0.75NA objective (Olympus) and projected onto the DAISY treated microfluidic chip to expose (500 mJ/cm$^2$) a circular area of 14 µm diameter. A custom software written in Python using the µManager API allowed the automatic exposure of DAISY in each compartment. After exposure, the chip was incubated with Biotin-NHS (Thermo Fisher Scientific, 5 mg ml$^{-1}$ in 250 mM borate buffer, pH 8.6) for 30 minutes, flushed with T250 buffer (10 mM Tris-HCl, pH 7.8 and 250 mM NaCl), and passivated with 0.1% Tween 20 in T250. Four chips were prepared in parallel and kept at room temperature (~20 °C) in a closed plastic box to prevent dehydration.

**Protein expression in the open system at the high gene density regime.**

The patterned and functionalized microfluidic chips were flushed with biotinylated Cy5 labeled circular DNA at a concentration of 50-75 nM and 70-105 nM streptavidin (ratio of 1:1.4) in T250 with 100 nM dummy DNA (non-coding DNA) and incubated for 1 h at room temperature. Afterwards, the chip was flushed with 2 ml of T250 buffer to remove any excess of DNA. The Cy5-DNA immobilization was verified by exciting the fluorophore with 12 mW at 647 nm (Colibri, Zeiss), collecting the emission signal with an 40x/0.75 NA objective (Olympus) through a Cy5 filter (Chroma), and imaging the signal on an EM-CCD camera (500 ms, 250 gain, iXon 987, Andor). Gene expression was performed on a standard optical microscope (Observer.Z1, Zeiss) with automated X, Y, and Z stage and auto-focus system. The temperature was controlled using a top stage heating chamber (Boldline, Okolab); the humidity was passively controlled by introducing small water reservoirs into the heating chamber. Images were acquired in an interval of 1.5 min with the same hardware settings as used for the DNA signal with an LED excitation intensity of 29.5 mW at 488 nm. The cell-free expression system was kept at 4 °C after preparation and introduced through a tubing (Tygon) using a 500 uL syringe pump (Gastight, Hamilton) with a constant negative flow rate of 0.4 uL/min (PHD Ultra, Harvard Apparatus).

**Computing the GFP level in individual compartments.**

To compute the GFP dynamics in the compartment, we outlined each compartment with a circular mask using Fiji and averaged the intensity. Since we found that PDMS fluoresced well within the GFP excitation spectrum and also bleached on a long time-scale (~hours), we subtracted the background signals taken from the vicinity of each compartment at every time point. As a final step and to further improve the sensitivity of our system, we subtracted the auto-fluorescence signal of the cell-free extract as measured inside a compartments without DNA for every time point.

**Protein expression in the open system at the low gene density.**

The functionalized microfluidic chips were flushed with biotinylated Cy5 labeled circular DNA at a concentration of 30 nM and 42 nM streptavidin (ratio of 1:1.4) in T250 and incubated for 30 minutes. The microfluidic chip was thoroughly washed with T250. Images were acquired on a custom-build single-molecule TIRF microscope. Illumination lasers with 488 nm (100 mW, OBIS, Coherent) and 647 nm (120 mW, OBIS, Coherent) were collinearly combined (DMSP605 and 5xBB1-E02, Thorlabs) through an objective mounted on a XYZ stage (MBT612D/M, Thorlabs) into a single-mode fiber (P5-460B-PCAPC-1, Thorlabs). The fiber output was coupled into a mirror collimator (RC08FC-P01, Thorlabs) to expand the laser diameter to 8 mm, guided through an achromatic lens (f=150 mm, AC254-150-A-ML, Thorlabs), re-directed by a mirror and filter cube (TRF59906, Chroma), and focused onto the back focal plane of the TIRF objective (60x, 1.49 NA, Nikon) in an angle to allow total internal reflection at the PDMS/glass and $H_2O$/glass interface. The cleaned emission signal was focused through a tube lens (TTL200, Thorlabs) onto an EM-CCD (iXon Ultra 888, Andor). The two lasers were selected with the Arduino microcontroller (35) and synchronized with the camera's exposure output trigger signal using digital modulation option of the laser controllers. The sample could be translated along XY (Märzhäuser Wetzlar) and the objective was mounted on a piezo stage to focus along Z (400 µm Fast PIFOC, PI) with an in-house build Delrin adapter for thermal insulation. The objective was further enclosed with resistive heating foil (HT10K, Thorlabs) to adjust the temperature using a PID controller (TE-48-20, TE Technology).

The temperature was adjusted and equilibrated for at least 15 min before the sample was placed on the holder. The DNA molecules were localized with 7 W/cm$^2$ at 647 nm (200 ms exposure time, 250 Gain). The microfluidic chip with the immobilized DNA was flushed with 10 µL of cell-free extract using a 10 µL pipette tip. The pipette tip was kept inside the inlet to maintain flow by gravity inside the main channel resulting in the replenishment of cell extract and removal of proteins that were produced from DNA immobilized in the main channel. Single-molecule expression dynamics were imaged with 15 W/cm$^2$ at 488 nm (200 ms exposure time, 250 gain) in an interval of 1 sec for a total acquisition time of 50-130 min.

We corrected the acquired movies for thermal drift with a cross-correlation algorithm (OpenCV) using the auto-fluorescent signal of the cell extract at 488 nm excitation. Then, individual compartments were outlined with the circular Hough transform (scikit-image). Inside the compartment boundaries, single molecules were localized with a local gradient algorithm (36) after correcting the uneven illumination of TIRF using a contrast limited adaptive histogram equalization (skimage). Intensity and background were extracted from the raw images. Trajectories were generated according to the following rules with trackpy: spots must not jump more than ~440 nm between consecutive frames (jump distance < 2 pixel) and stayed there for at least 3 sec with a memory of 1 sec to account for blinking events.

**Calculation of autocorrelation functions of production rates.**

The autocorrelation function is computed for each compartment from the production rate $x(t)$ after 20 minutes with the time-average production rate $\langle x \rangle$, variance $\sigma_x^2$ for a time delay $\tau$ as (Fig. S12, Fig. S14, and Fig. S19):

$$A(\tau) = \frac{\langle (x(t) - \langle x \rangle)(x(t+\tau) - \langle x \rangle \rangle}{\sigma_x^2}$$

Since individual compartments show high variability in the time-averaged production rates, we also computed the ensemble-averaged by averaging individual ACFs (Fig. S17 and Fig. S19).

**Theoretical models of CI regulation.**

An equilibrium model can be derived according to the mass-action law for simple

$$I + D \leftrightharpoons ID$$

and cooperative binding

$$I + D \leftrightharpoons ID$$
$$I + ID \leftrightharpoons I_2D$$

of inhibitor *I* binding to its operator site *D*. They give an equilibrium dissociation constants according to $K_D = I^n D/I_n D$ with *n* = 1,2,... . By assuming no self-inhibition (here only valid for short times), the amount of *I* during protein expression is given by

$$I = k_{P,I}\, t$$

where $k_{P,I}$ (nM/min) is the production rate of the repressor in the cell extract. Together with the equilibrium equations, the amount of free DNA with $D = D_0 - ID$, and the reporter production with $dg/dt = k_{P,G} D$, with $k_{P,G}$ (1/min) being the production rate of the reporter, the final form is

$$\frac{dg}{dt} = \frac{k_{P,G}\, D_0}{\frac{(k_{P,I}\, t)^n}{K_D} + 1}$$

Integration with non-cooperative binding (*n* = 1) from *g = 0* to *G* and t = 0 to *T,* gives the accumulation of reporter *G* according to

$$G = \frac{k_{P,G}\, D_0 K_D}{k_{P,I}} \ln \frac{k_{P,I}\, T + K_D}{K_D}$$

with $K_D$ (nM) being the affinity constant.

Cooperative binding (*n* = 2) of inhibitor *I* changes the course of the fluorescent reporter *G* with

$$G = \frac{k_{P,G}\, D_0}{\sqrt{k_{P,I}^2/K_D}} \tan^{-1} \sqrt{\frac{k_{P,I}^2}{K_D}}\, T$$

where $K_D$ (nM$^2$) is the affinity constant. Comparing the experimental data with the derived models (Fig. S16), gives an estimate of around a 100-fold stronger affinity of CI to the operators for the production of *G*.

The stochastic simulations to estimate the $k_{on}$ rates for CI to the operator sites were conducted with COPASI using the stochastic integration by the direct method (Gillespie algorithm). The following equations were implemented inspired by the model from Isaacs *et al.* (19) with the parameters given in Table S2. Equilibrium binding of protein *P* to the three operator sites on DNA *D*:

$$P + D \leftrightharpoons D_1,\ P + D_1 \leftrightharpoons D_2,\ P + D_2 \leftrightharpoons D_3$$

Transcription reactions with the basal (*$TX_0$*) and activated (*$TX_a$*) states to produce mRNA *R*:

$$D \xrightarrow{TX_0} R,\ D_1 \xrightarrow{TX_a} D_1 + R,\ D_2 \xrightarrow{TX_a} D_2 + R,$$

Translation reaction with translation rate *TL*:

$$R \xrightarrow{TL} R + P,$$

Degradation and inactivation reactions are defined for mRNA with the rate of *mdeg* and all other rates with the temperature dependent rates (Table S3):

$$R \xrightarrow{mdeg} \emptyset,$$

$$P \rightarrow \emptyset,\ D_1 \rightarrow D,\ D_2 \rightarrow D_1,\ D_3 \rightarrow D_2$$

The data plotted in Fig. 3D is the number of proteins *P* produced in an interval size of 15 seconds and a total simulation time of 60 min (ensemble averaged production rates was also extracted only after 20 min in steady-state). Simulations were run from 31 to 41 °C in 1 °C steps and each temperature was simulated for N=20.

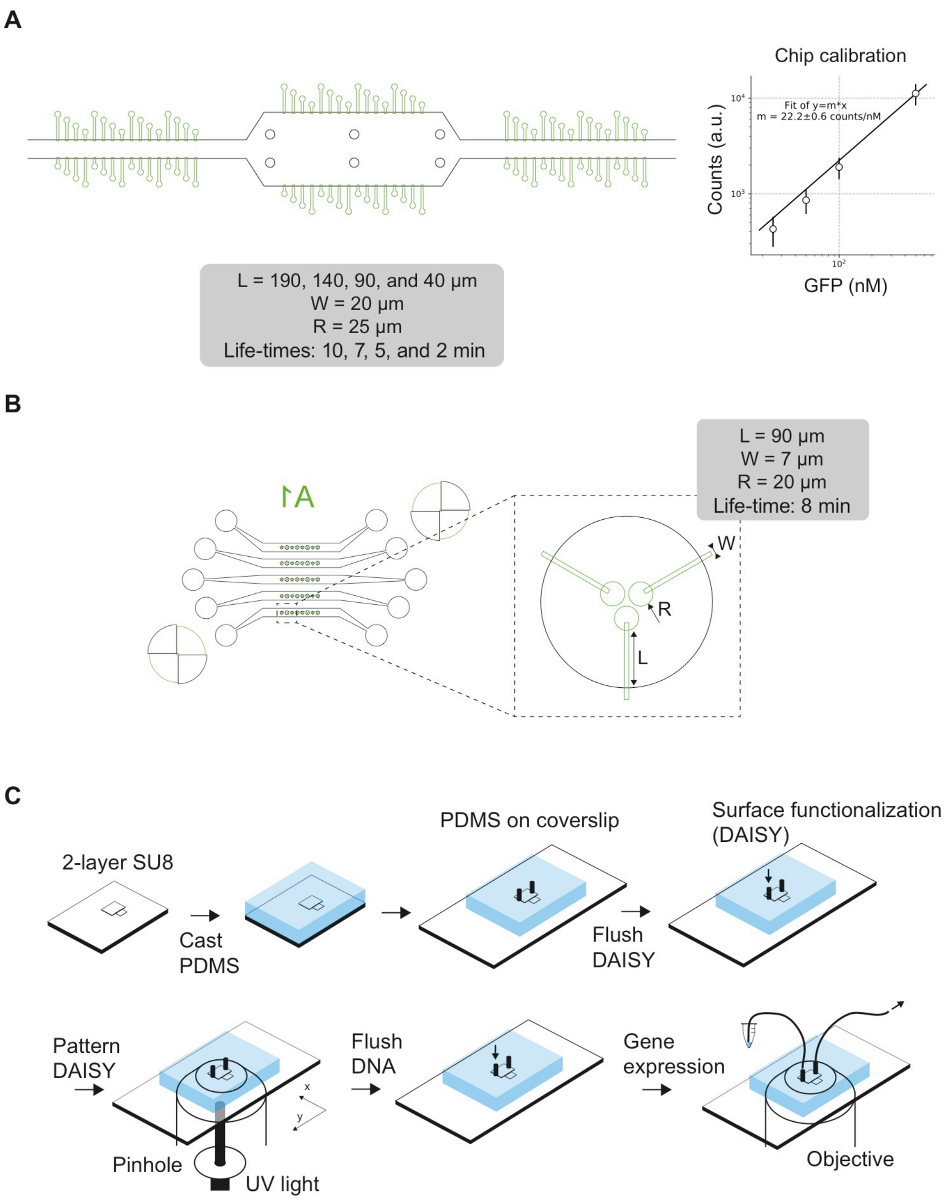


**Fig. S1. Design and assembly of microfluidics chips.** (**A**) Chip design for high molecule numbers in the high-density regime. The GFP concentration was calibrated by flushing known concentrations of purified GFP in 1x PBS. (**B**) Chip design for single-molecule experiments on TIRF microscope. 5 lanes to perform multiple experiments at different conditions. (**C**) PDMS chips are fabricated with 2-layer SU8 structures, flushing of chemicals (Methods) directly onto the chip, and *in situ* DNA patterning and processing until gene expression is performed with *E. coli* cell extract.

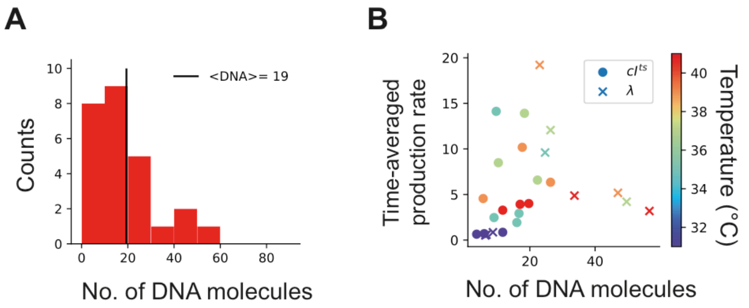


**Fig. S2. Numbers of DNA molecules and time-average production rates during experiments at the low-density regime.** (**A**) Histogram of detected DNA spots in individual compartments. (**B**) Time-averaged production rates against the number of DNA spots (with bistable "λ" and monostable "cI$^{ts}$" GRN) in individual compartments at various temperatures (color-coded).

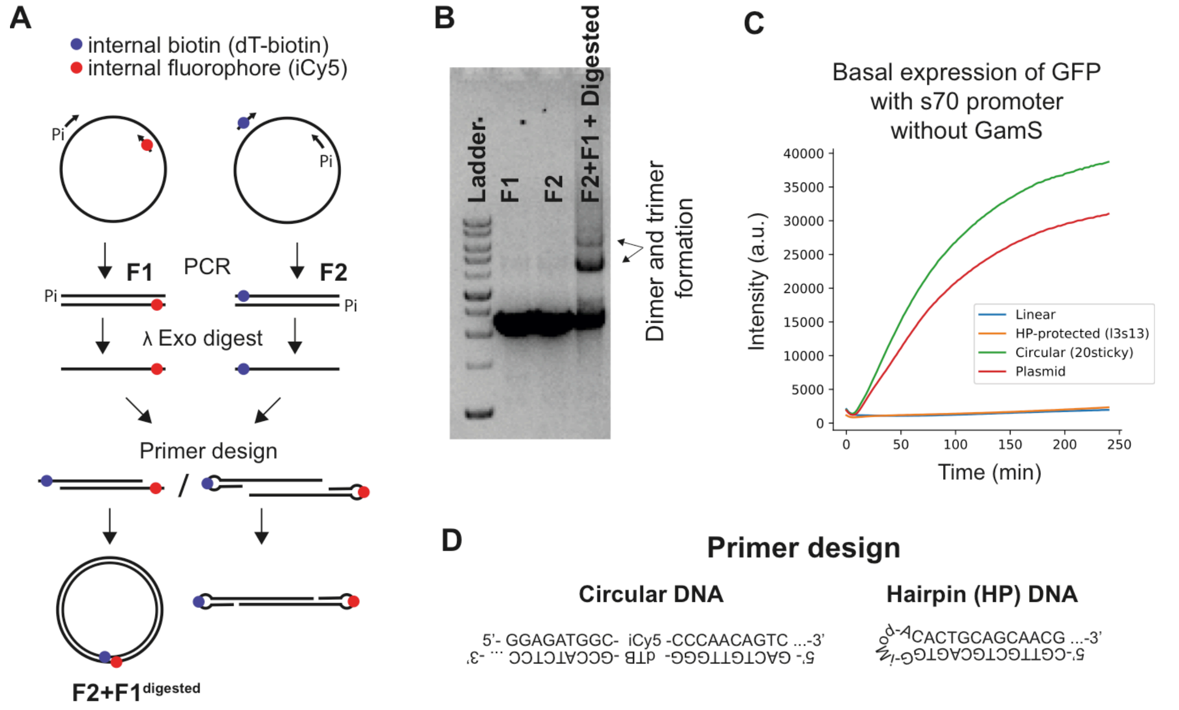


**Fig. S3. DNA protection by DNA circularization.** (**A**) 2-step protocol to modify the ends of double-stranded DNA fragments with PCR. Step 1: Two separate PCR reactions are started with two primer pairs (Methods, Table S1) on the same plasmid. Step 2: Digestion of double-stranded PCR fragments gave single-stranded DNA fragments. The selective digestion of one strand by lambda exonuclease was controlled with phosphorylated primers. The two single-stranded DNA fragments annealed simultaneously to give a double-stranded fragment with end modifications, e.g. circular DNA or hairpin structure. (**B**) Agarose gel with ladder (1kb DNA ladder, ThermoFisher Scientific), F1, F2, and digested F1+F2 fragments (from left to right). The circular DNA could be identified after annealing F1 and F2. Higher bands (dimers and trimers) are formed through self-binding at the high concentrations. (**C**) GFP production in solution experiments from different DNA sources ("Linear" = linear dsDNA as negative control; "HP-protected (l3s13") = Hairpin structure with 3 and 13 bases in loop and stem region, respectively; "Circular (20sticky)" = circular DNA with 19 (=20 - 1 modification) bases of single-stranded overhang; "Plasmid" = plasmid as positive control) in *E. coli* cell extract without GamS at 32 °C (upper right panel). (**D**) Overlapping sequences of primers for circular and hairpin modifications.

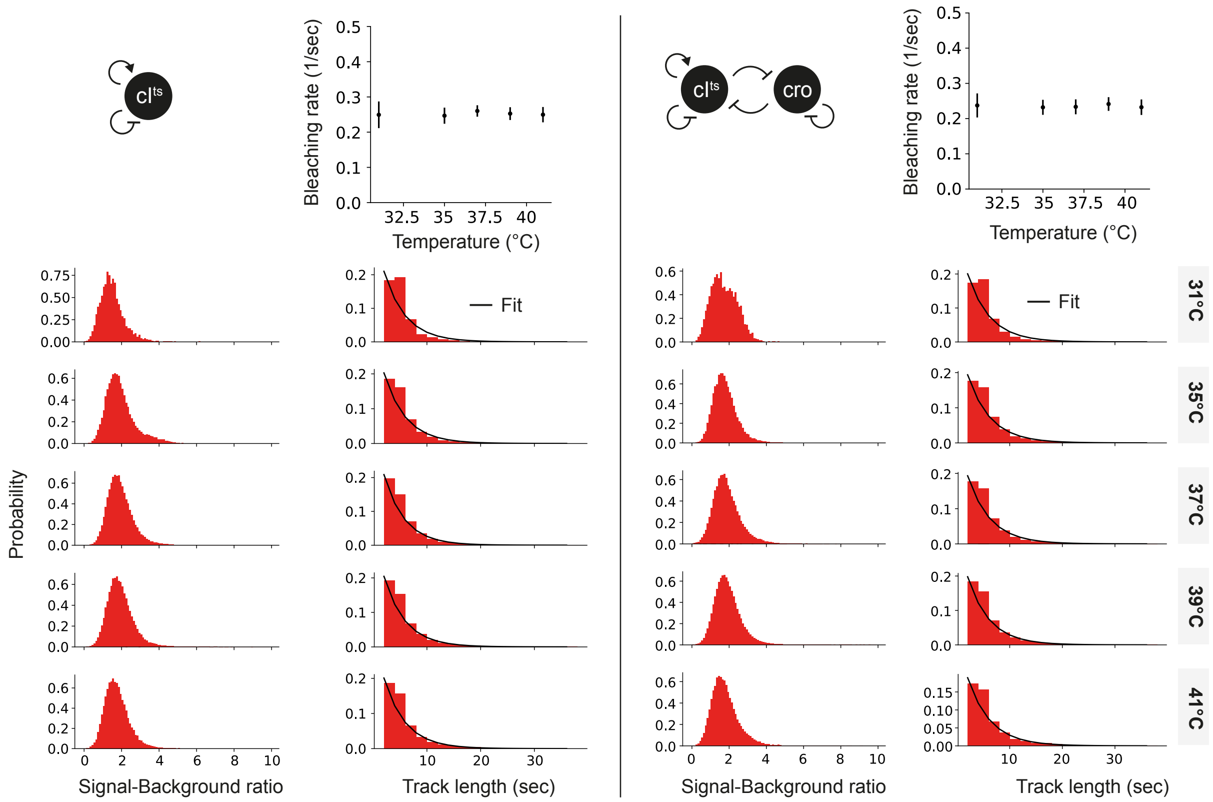


**Fig. S4. Properties of individual cI$^{ts}$-mVenus proteins.** Signal-to-background ratios of single cI$^{ts}$-mVenus at the various temperatures (left column for both GRNs, separated by a solid line). The time from the first encounter of a single mVenus spot till it vanished is computed for all spots, pooled and plotted as track length in the right column for the various temperatures (left column for both GRNs, separated by a solid line). The probability was fitted with a mono-exponential decay and gave values plotted against temperature in the right uppermost figure for each GRN.

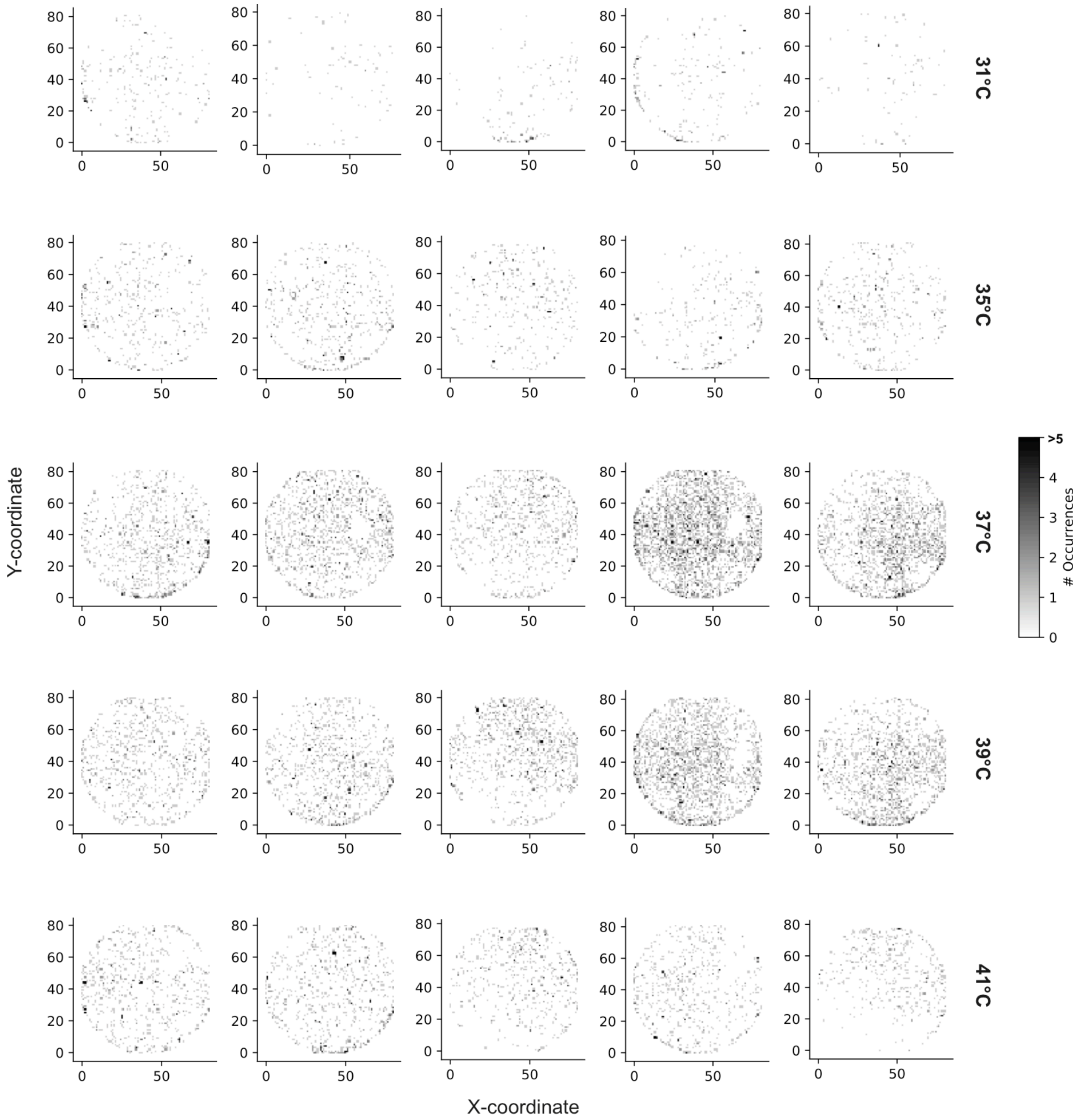


**Fig. S5. Spatial distribution and clustering of single CI$^{ts}$-mVenus proteins.** 2-D spatial histogram of spot counts integrated over the entire course of the experiment in 5 representative compartments at the indicated temperatures. Particles appeared evenly distributed at >37 °C due to unspecific adsorption of denatured and therefore sticky CI$^{ts}$-mVenus and non-optimal solution conditions. For other *in vitro* single-molecule studies, buffer and surface conditions can be readily changed towards minimizing unspecific adsorption (37), that was however not feasible due to the current working conditions of the cell-free extract.

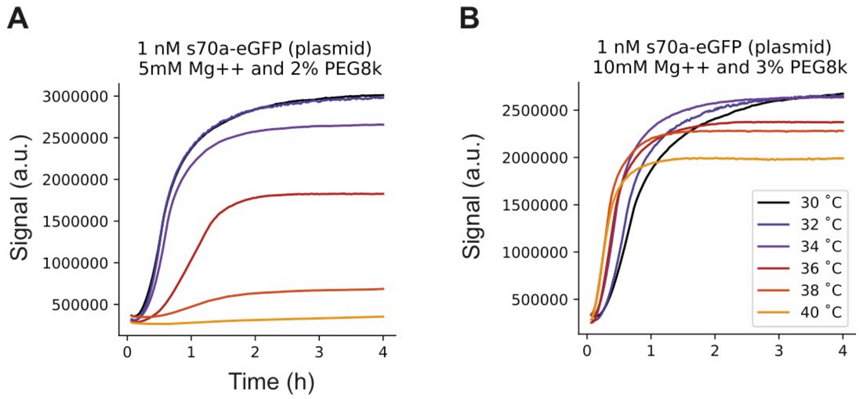


**Fig. S6. Optimization of expression conditions for the cell-free extract.** (**A**) Fluorescent signal of GFP expression in solution experiments from plasmid at 1 nM concentration at various constant temperatures (color-coded) and 5 mM $Mg^{2+}$ and 2% PEG8000 (crowding agent). (**B**) Fluorescent signal of GFP expression in solution experiments from plasmid at 1 nM concentration at various constant temperatures (color-coded) and 10 mM $Mg^{2+}$ and 3% PEG8000 (crowding agent). A higher concentration of $Mg^{2+}$ and PEG8000 improved expression and led to similar amounts of GFP within the temperature range.

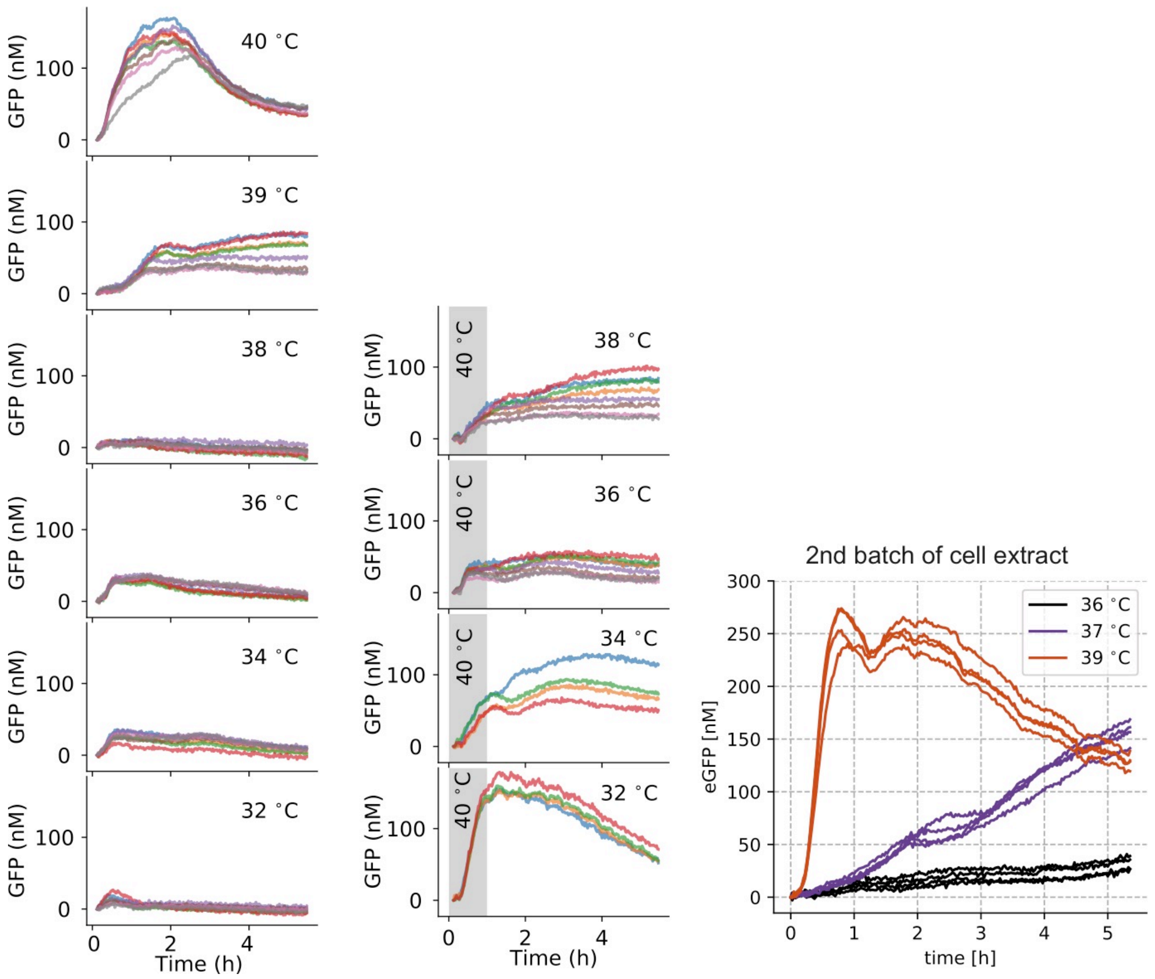


**Fig. S7. Response of the bistable GRN in the high-density regime.** The fluorescent signal of GFP under the $P_R$ promoter in single compartments and various temperatures (first column). The bistability of the GRN was tested with initial protein expression at 40 °C (gray region in second column) and a drop in temperature (indicated on the right side in each plot of the second column). The GFP response was also obtained with a 2nd batch of cell-extract (third column).

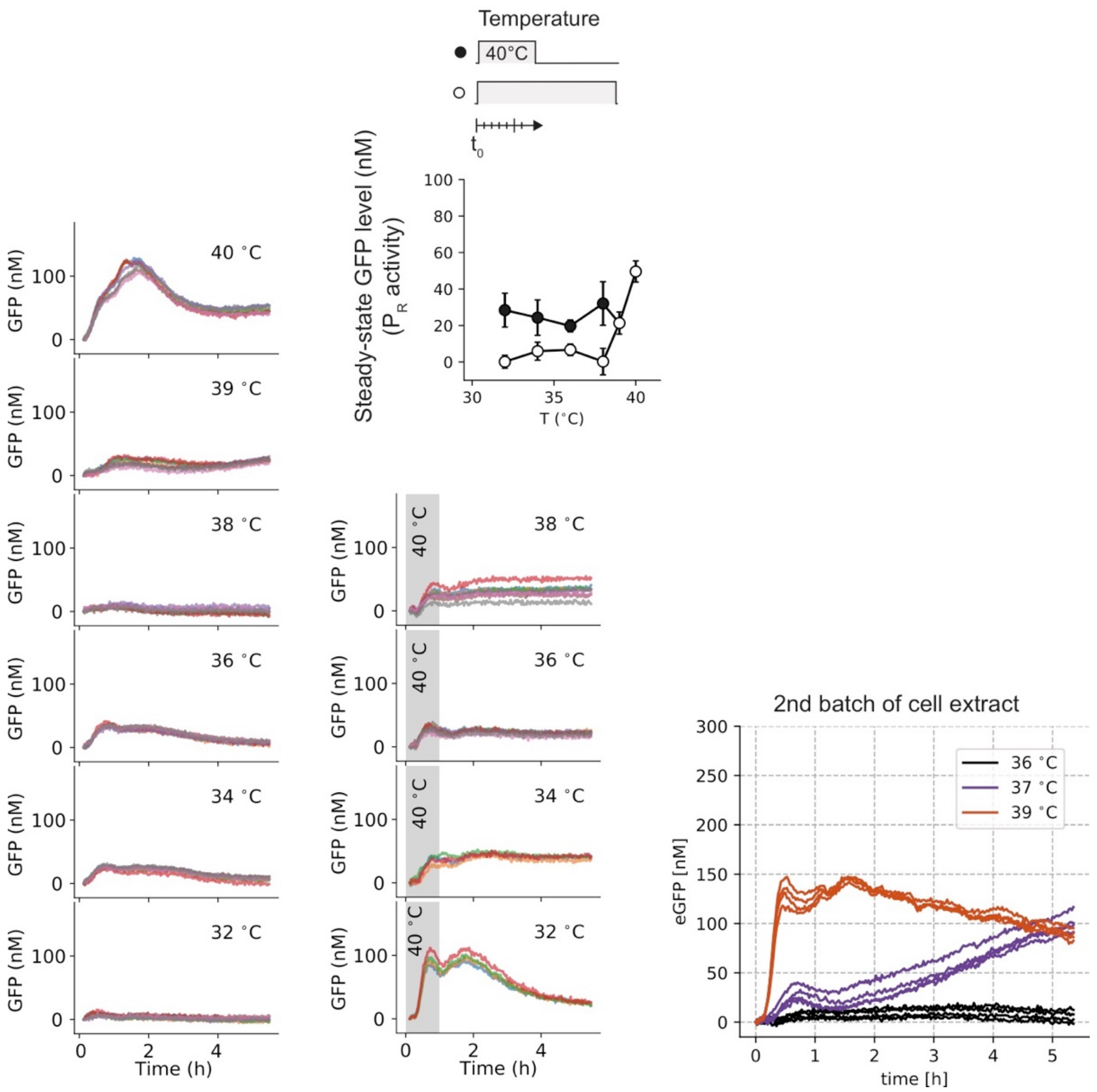


**Fig. S8. Response of the bistable GRN in the high-density regime at shorter compartment life-times.** The fluorescent signal of GFP under the $P_R$ promoter in single compartments with 7 min life-time (see Fig. S1A) and various temperatures (first column). The bistability of the GRN was tested with initial protein expression at 40 °C (gray region in second column) and a drop in temperature (indicated on the right side in the subfigures of the second column). Steady-state values are plotted for the various temperature inputs (uppermost figure in second column). The GFP response was also obtained with a 2nd batch of cell-extract. (third column).

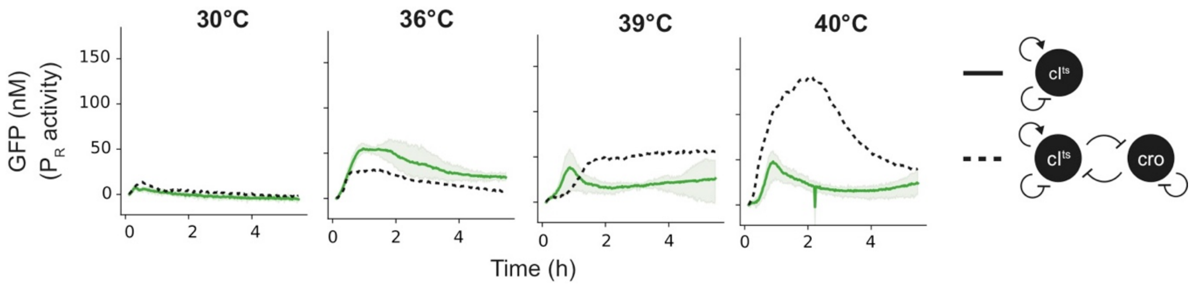


**Fig. S9. Response of the monostable GRN in the high-density regime.** The fluorescence signal of GFP under the $P_R$ promoter without the *cro* gene in single compartments and various temperatures. The GFP dynamics of the bistable GRN with the *cro* gene are indicated as dashed line for comparison.

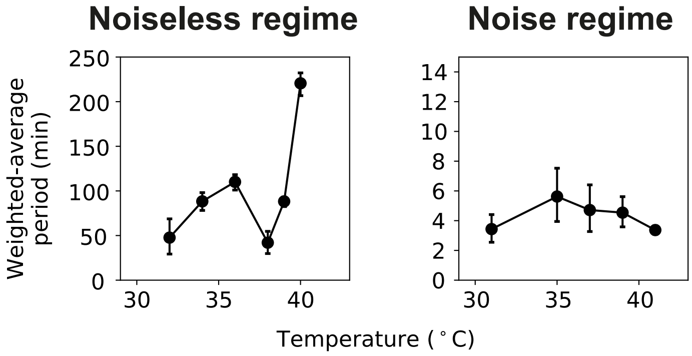


**Fig. S10. Characteristic time scale of computation in the bistable GRN.** A fast Fourier transform (FFT) of the full expression dynamics (with onset time) in the high gene density (left) and low gene density (right) regimes was used to compute the frequency domain. The frequencies were then averaged with the weights of the corresponding FFT power spectrum (excluding the zero and noise frequencies, $0 < f_i < 1$/min) and plotted for all temperatures. Whereas the noiseless regime showed dynamics on a time scale of ~1 h, the dynamics in the noise regime happened on the ~5 min time scale.

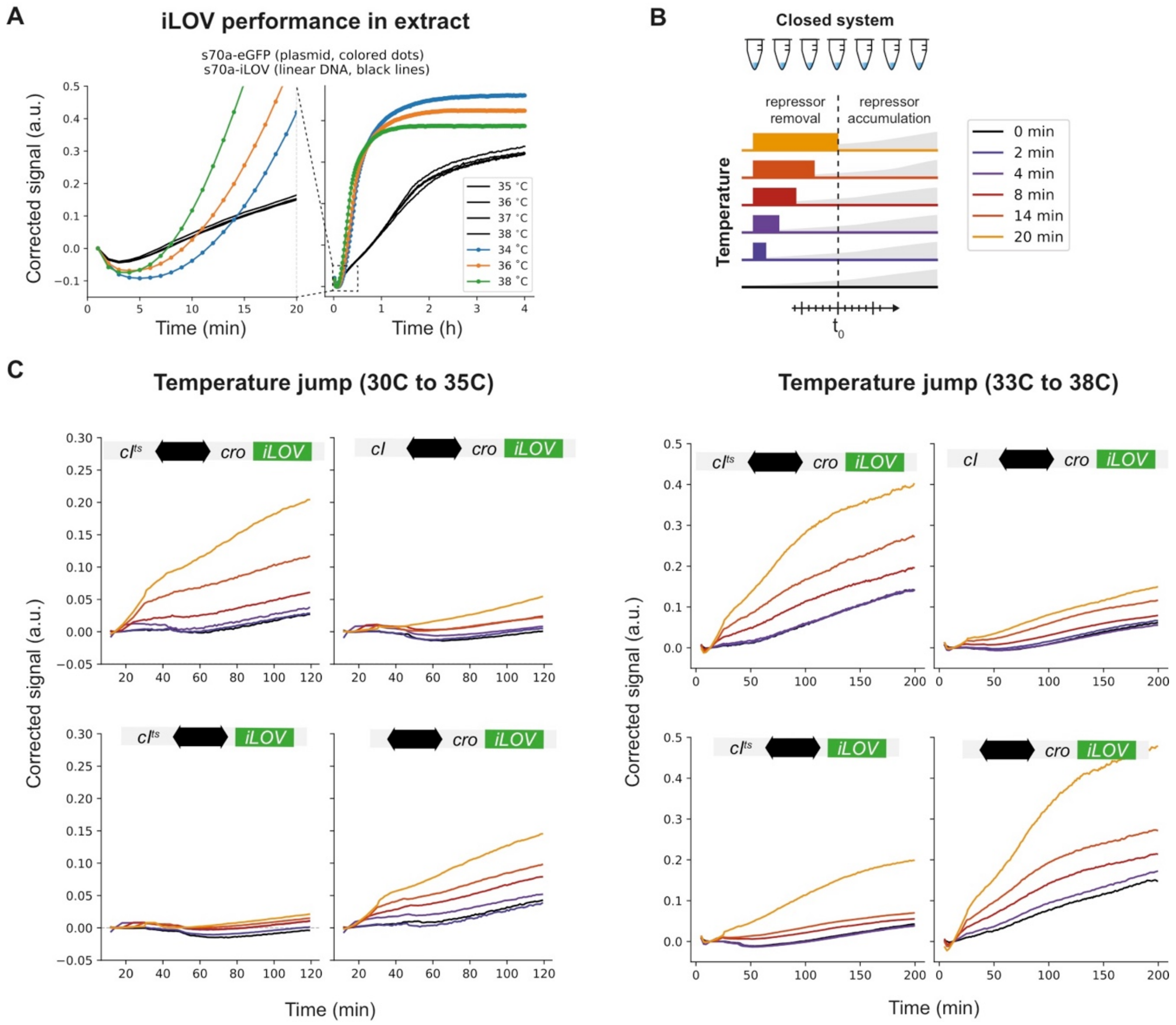


**Fig. S11. Expression dynamics of the GRNs in solution experiments.** (**A**) The production dynamics of iLOV against the dynamics of GFP production in cell extract at various temperatures under a constitutive promoter. (**B**) The protocol to verify the decision-making of the GRNs in solution. The initial temperature jump deactivated temperature-sensitive $CI^{ts}$ and allows to accumulate after the drop in temperature. (**C**) Temperature jumps from 30/33 °C (left/right) to 35/38 °C (left/right) for the indicated GRNs (upper row: the bistable GRN with and without the temperature sensitive mutation, lower row: the monostable GRNs with the temperature-sensitive $cI^{ts}$ and with cro only). While the bistable wild-type GRN showed little temperature dependence compared to the bistable GRN due to the missing temperature mutation, the monostable GRN without the *cro* gene is not dependent on temperature since the GRN can return to their initial promoter activity.

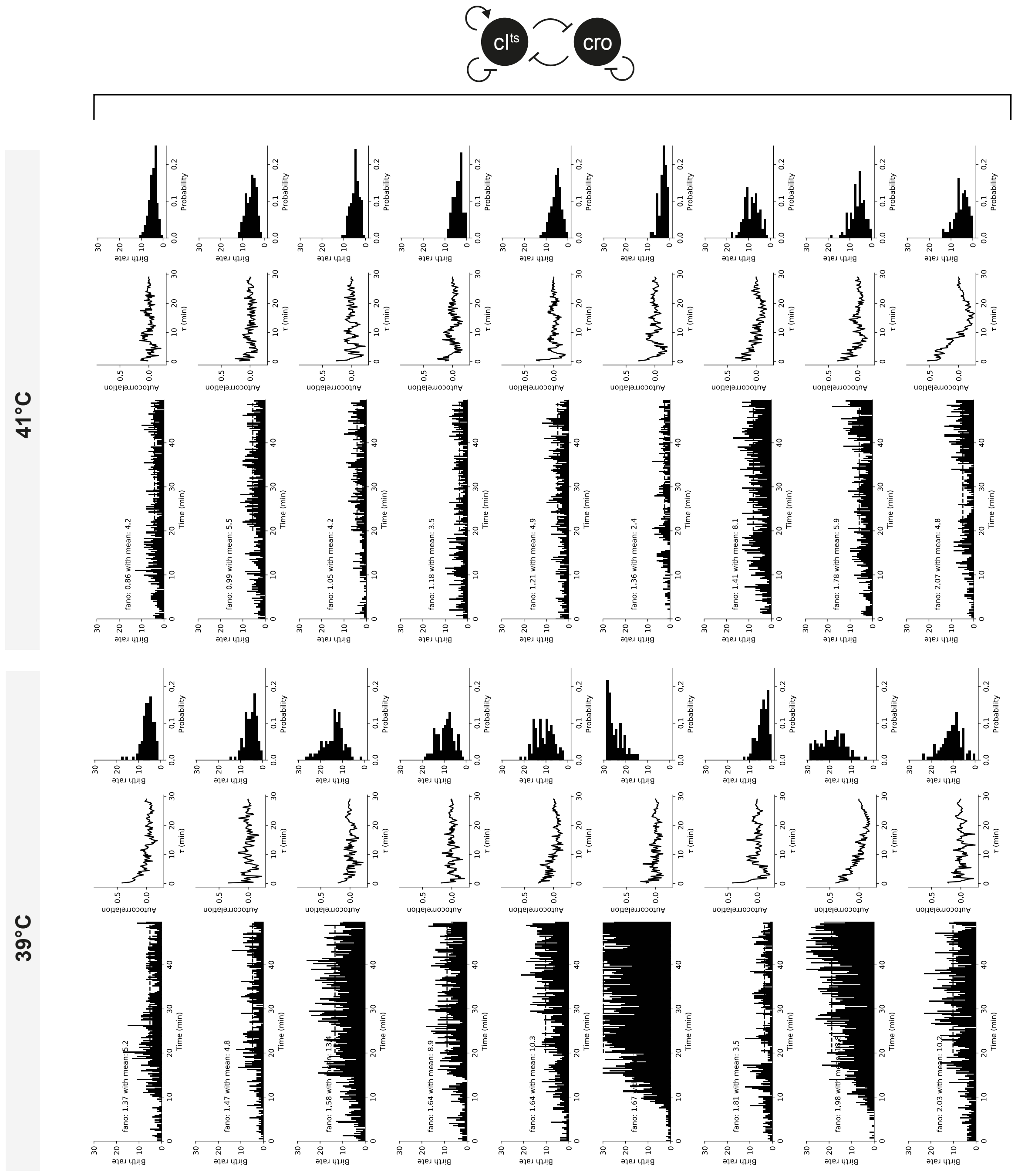
cI^ts
cro
41°C
39°C
fano: 0.86 with mean: 4.2
fano: 0.99 with mean: 5.5
fano: 1.05 with mean: 4.2
fano: 1.18 with mean: 3.5
fano: 1.21 with mean: 4.9
fano: 1.36 with mean: 2.4
fano: 1.41 with mean: 8.1
fano: 1.78 with mean: 5.9
fano: 2.07 with mean: 4.8
fano: 1.37 with mean: 5.2
fano: 1.47 with mean: 4.8
fano: 1.64 with mean: 8.9
fano: 1.64 with mean: 10.3
fano: 1.81 with mean: 3.5
fano: 2.03 with mean: 10.2
Birth rate
Time (min)
Autocorrelation
τ (min)
Probability

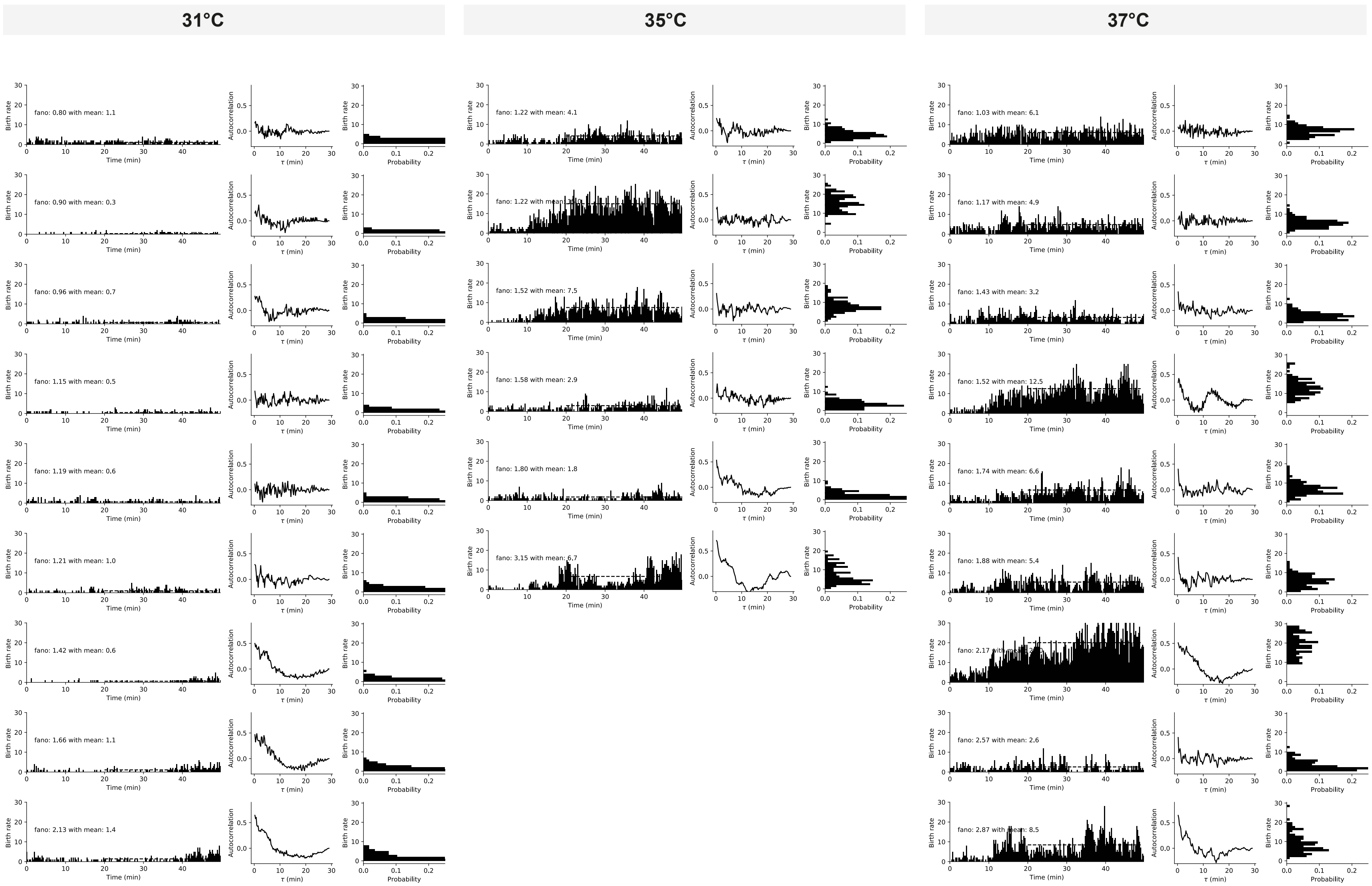
31°C
35°C
37°C
fano: 0.80 with mean: 1.1
fano: 0.90 with mean: 0.3
fano: 0.96 with mean: 0.7
fano: 1.15 with mean: 0.5
fano: 1.19 with mean: 0.6
fano: 1.21 with mean: 1.0
fano: 1.42 with mean: 0.6
fano: 1.66 with mean: 1.1
fano: 2.13 with mean: 1.4
fano: 1.22 with mean: 4.1
fano: 1.22 with mean
fano: 1.52 with mean: 7.5
fano: 1.58 with mean: 2.9
fano: 1.80 with mean: 1.8
fano: 3.15 with mean: 6.7
fano: 1.03 with mean: 6.1
fano: 1.17 with mean: 4.9
fano: 1.43 with mean: 3.2
fano: 1.52 with mean: 12.5
fano: 1.74 with mean: 6.6
fano: 1.88 with mean: 5.4
fano: 2.17 with
fano: 2.57 with mean: 2.6
fano: 2.87 with mean: 8.5
Birth rate
Time (min)
Autocorrelation
τ (min)
Probability

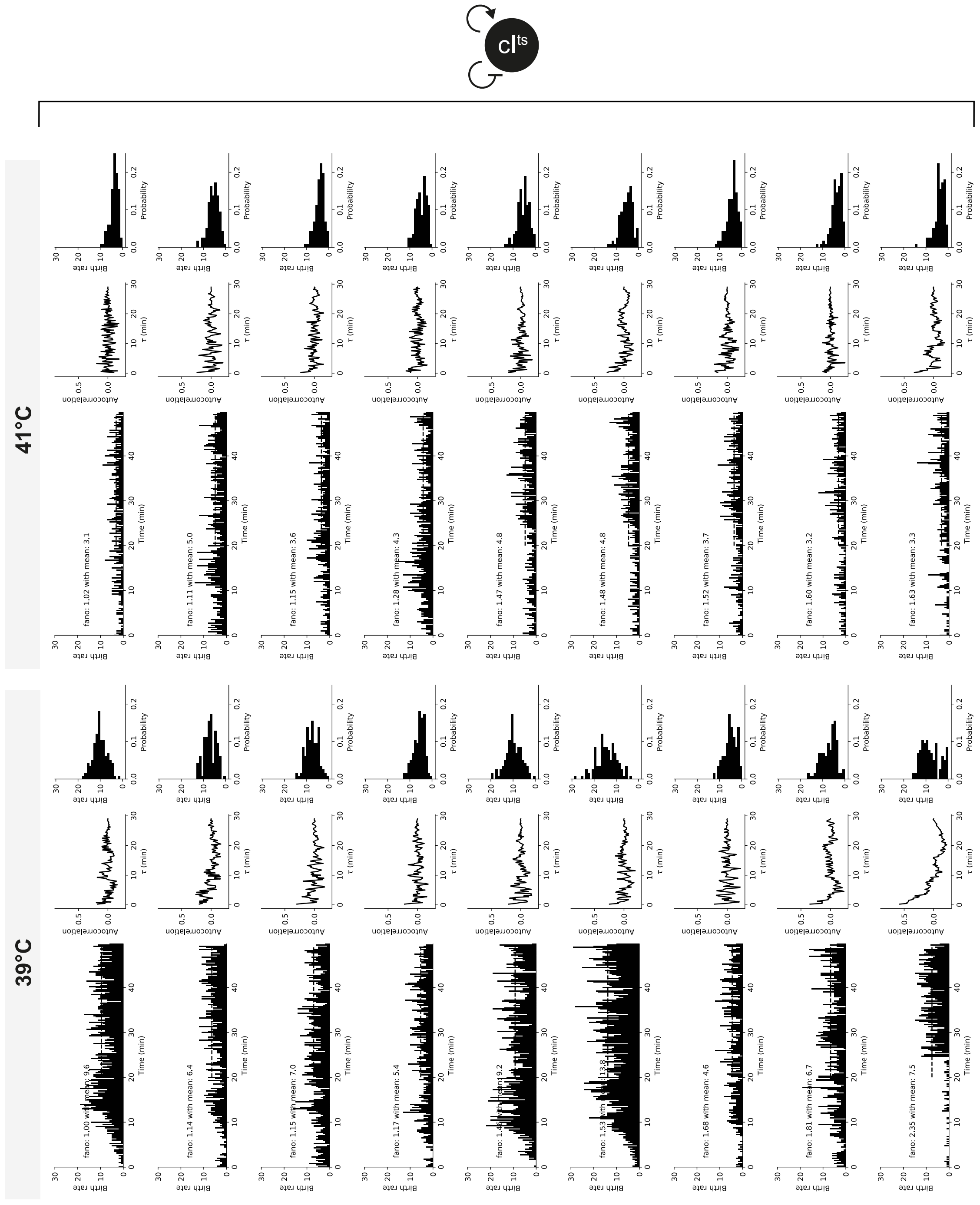
39°C
41°C

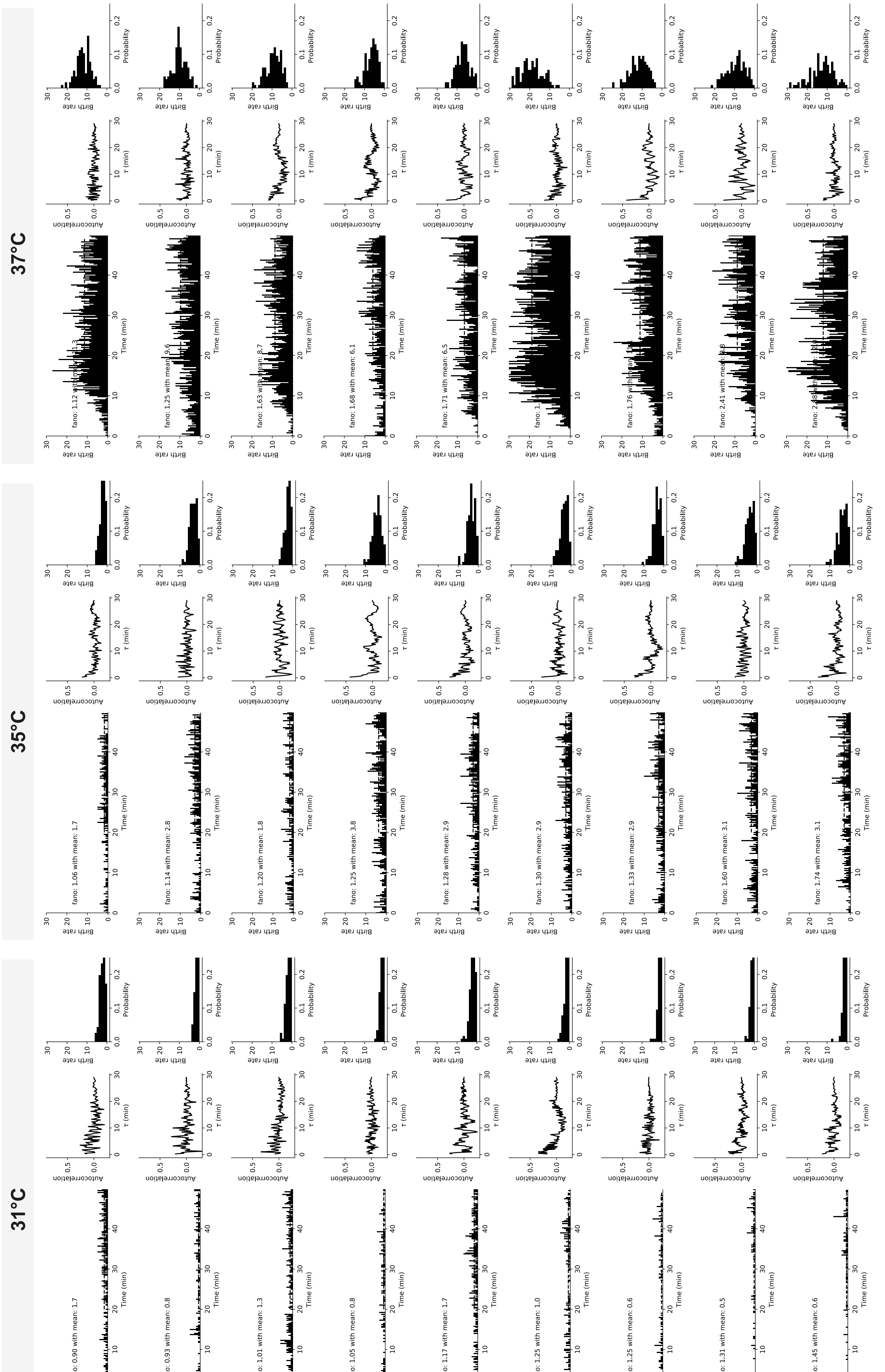
31°C
35°C
37°C
fano: 0.90 with mean: 1.7
fano: 0.93 with mean: 0.8
fano: 1.01 with mean: 1.3
fano: 1.05 with mean: 0.8
fano: 1.17 with mean: 1.7
fano: 1.25 with mean: 1.0
fano: 1.25 with mean: 0.6
fano: 1.31 with mean: 0.5
fano: 1.45 with mean: 0.6
fano: 1.06 with mean: 1.7
fano: 1.14 with mean: 2.8
fano: 1.20 with mean: 1.8
fano: 1.25 with mean: 3.8
fano: 1.28 with mean: 2.9
fano: 1.30 with mean: 2.9
fano: 1.33 with mean: 2.9
fano: 1.60 with mean: 3.1
fano: 1.74 with mean: 3.1
fano: 1.68 with mean: 6.1
fano: 1.71 with mean: 6.5
Birth rate
Time (min)
Autocorrelation
τ (min)
Probability

**Fig. S12. Examples of single compartments with production rates at different temperatures and GRNs.** The time-averaged production rates (shown in subfigures), Fano factor (shown in subfigures), ACFs and probability distributions are computed from the region as indicated by the dashed line. Total number of experiments for the bistable (upper two pages) and monostable (lower two pages) GRN in the order from 31 to 41 °C were 9, 6, 9, 18, 9 and 9, 12, 9, 9, 9, respectively.

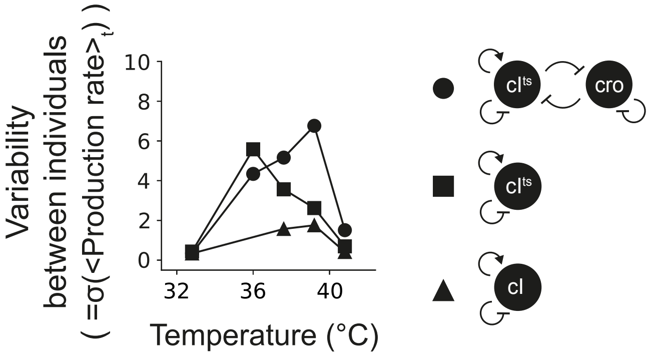


**Fig. S13. Variability between individual cell models.** The variability was computed as standard deviation from the time-averaged production rates of individual compartments for the various temperatures and indicated GRNs.

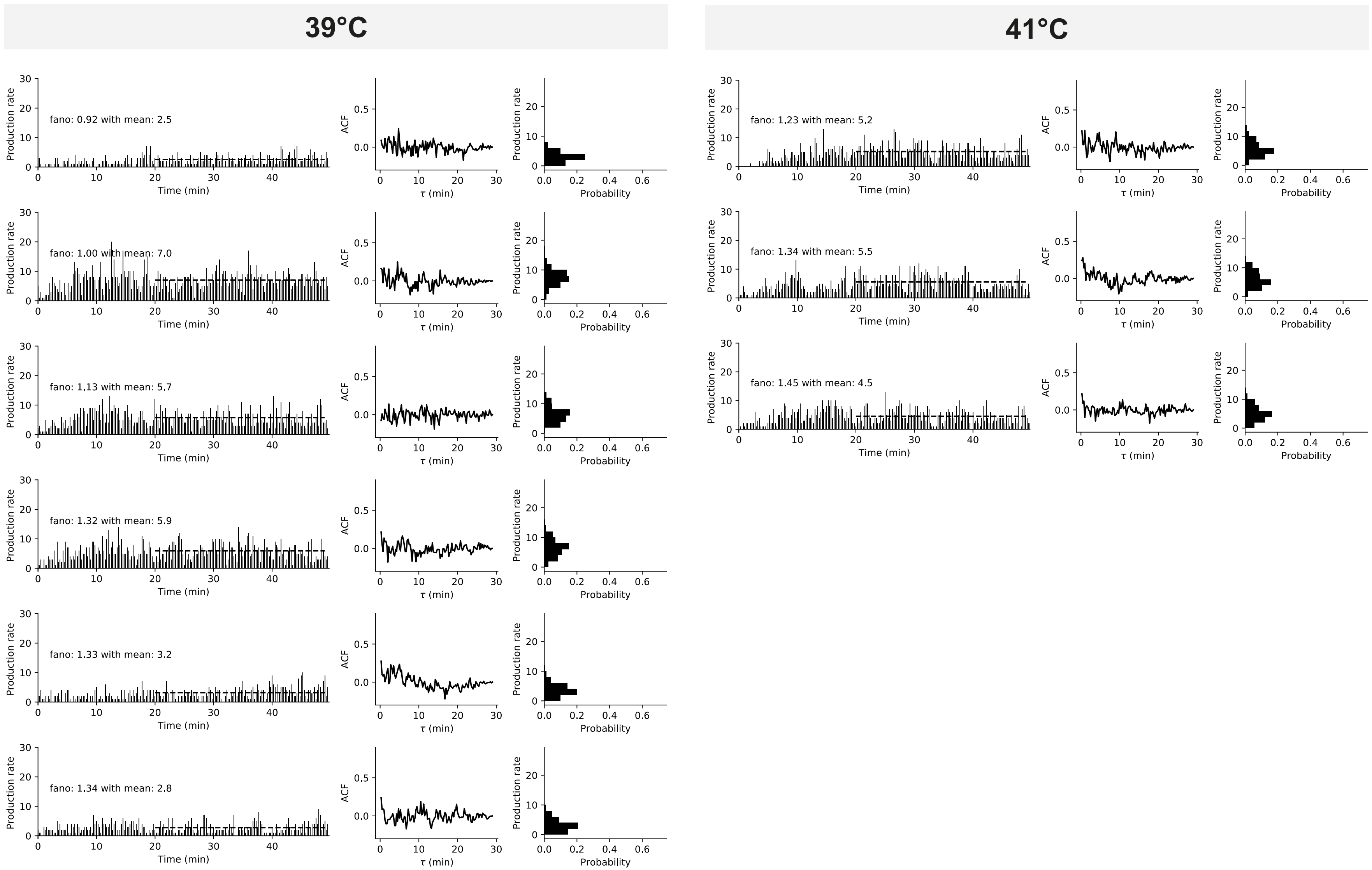
39°C
41°C
fano: 0.92 with mean: 2.5
fano: 1.00 with mean: 7.0
fano: 1.13 with mean: 5.7
fano: 1.32 with mean: 5.9
fano: 1.33 with mean: 3.2
fano: 1.34 with mean: 2.8
fano: 1.23 with mean: 5.2
fano: 1.34 with mean: 5.5
fano: 1.45 with mean: 4.5
Production rate
Time (min)
ACF
τ (min)
Probability

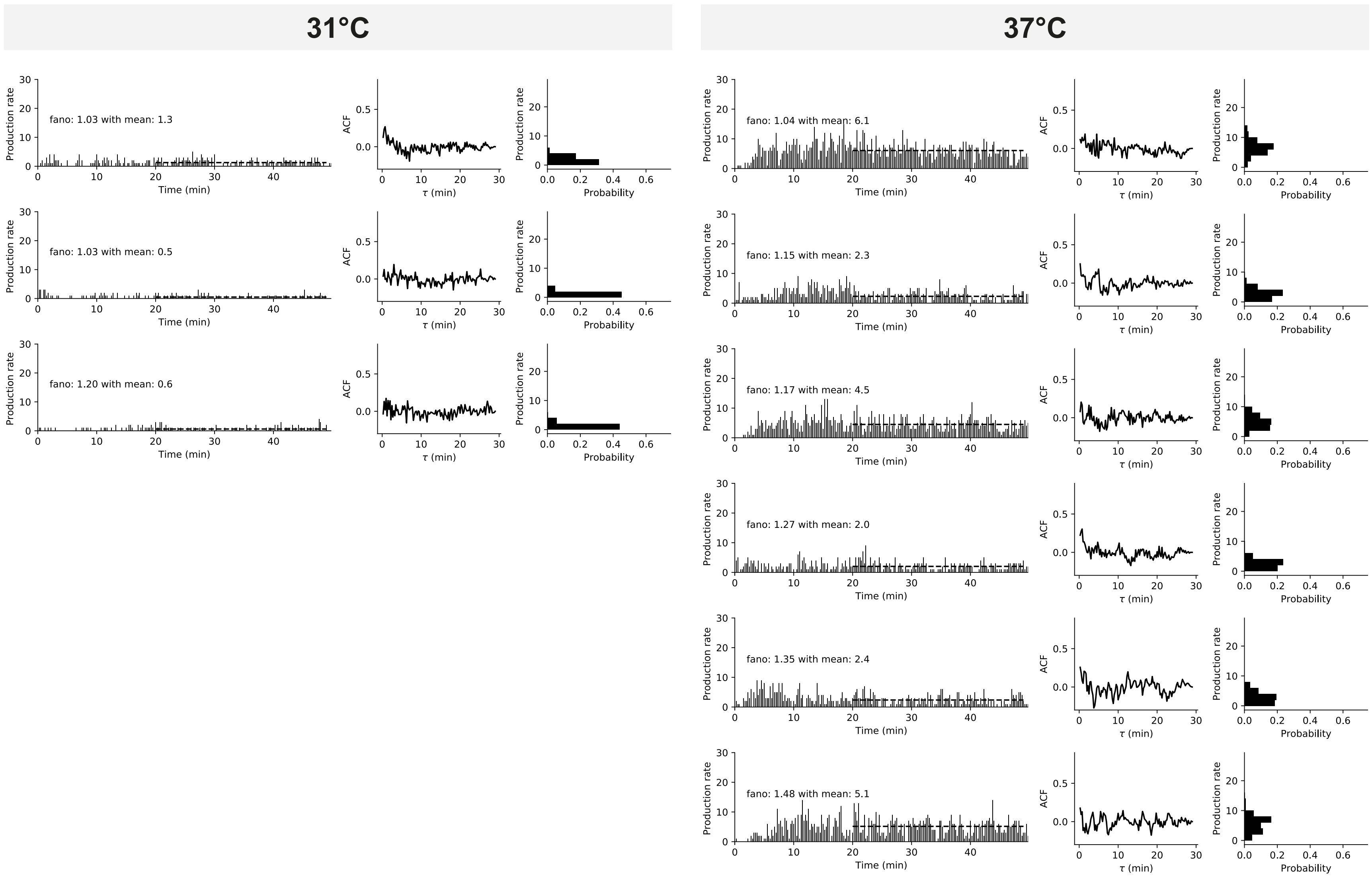
31°C
37°C
fano: 1.03 with mean: 1.3
fano: 1.03 with mean: 0.5
fano: 1.20 with mean: 0.6
fano: 1.04 with mean: 6.1
fano: 1.15 with mean: 2.3
fano: 1.17 with mean: 4.5
fano: 1.27 with mean: 2.0
fano: 1.35 with mean: 2.4
fano: 1.48 with mean: 5.1
Production rate
Time (min)
ACF
τ (min)
Probability

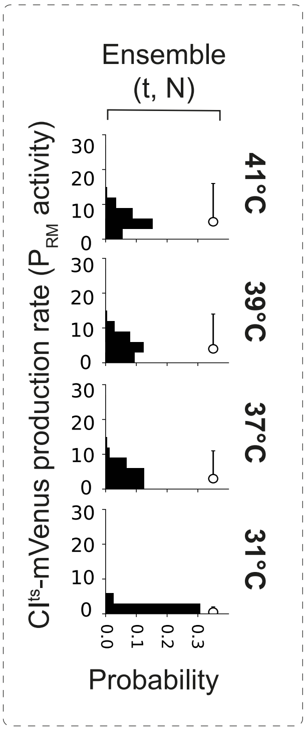


**Fig. S14. Single compartments with production rates at different temperatures with the wild-type monostable GRN.** The time-averaged production rates (shown in subfigures), Fano factor (shown in subfigures), ACFs and probability distributions are computed from the region as indicated by the dashed line. (left column). The ensemble production rate as in Fig. 2C and Fig. 3A in the main text (dashed box). Total number of experiments for the wild-type monostable GRN in the order from 31 to 41 °C were 3, 6, 6, 3.

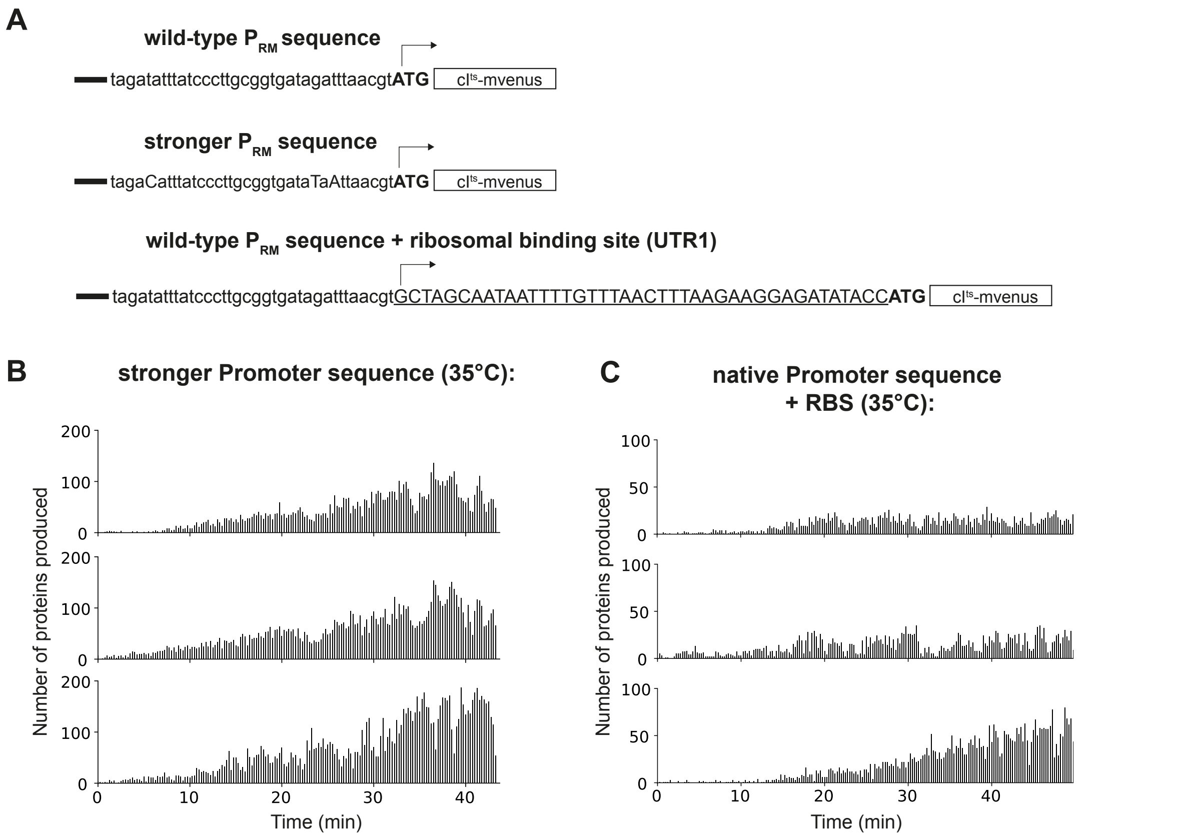


**Fig. S15. Production rates in single compartments with change of transcription and translation elements in the monostable GRN without cro gene.** (**A**) Wild-type sequence of $P_{RM,}$ a consensus promoter for $\sigma_{70}$, and the native promoter with ribosomal binding site at the 5'-UTR of *cI^ts^*. (**B**) Proteins produced with the consensus promoter sequence. The signal was measured as in Fig. 1E, Fig. 2C, Fig. 3A, and Fig. 4C. (**C**) Proteins produced with the RBS and native $P_{RM}$. The signal was measured as in Fig. 1E, Fig. 2C, Fig. 3A, and Fig. 4C.

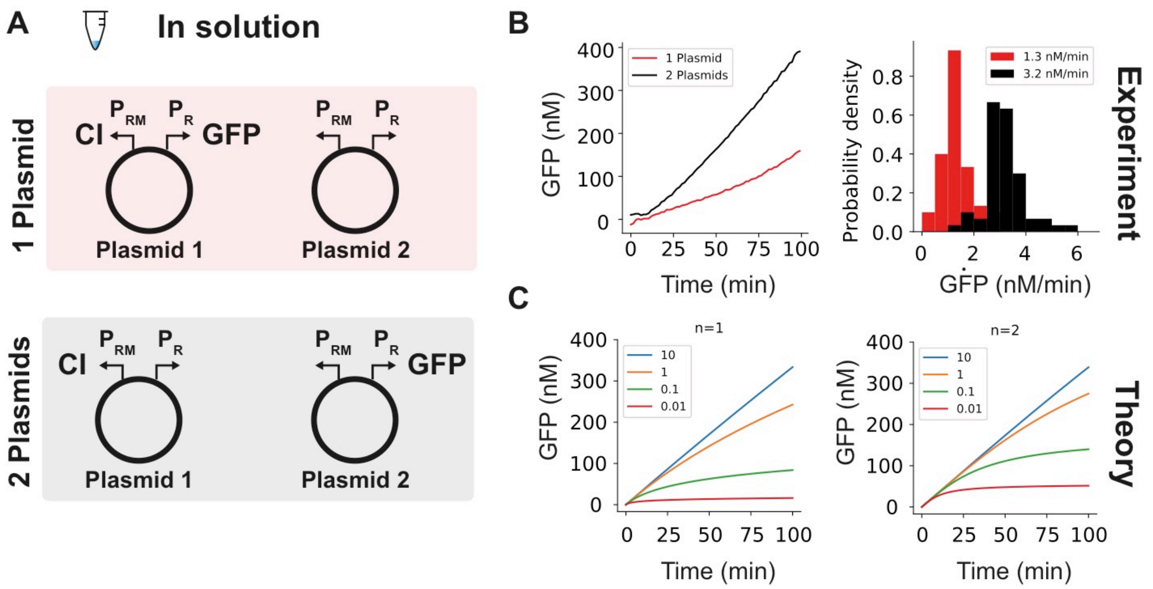


**Fig. S16. Solution experiment with co-expression of CI$^{ts}$ and GFP.** (**A**) First set (red region): The first plasmid encoded both CI and GFP with a promoter architecture as in the bistable GRN. The other plasmid encoded no gene, but leveled the total DNA concentration. Second set (black region): The first plasmid only encoded CI under the native $P_{RM}$ promoter architecture. The other plasmid encoded GFP under the $P_R$ promoter with the native promoter architecture. (**B**) Protein expression was performed in cell extract at 32 °C (no deactivation) with two different pairs of plasmids. Increase of GFP signal over time is shown (first plasmid set = red, second plasmid set = black). The data was clipped to the linear expression regime (the dynamics of closed systems eventually slow down due to energy consumption). The rate of GFP production is plotted with the average rates indicated in the figure legend (right plot). (**C**) Theoretical dynamics of equilibrium binding of CI to DNA (Methods). Comparing the experimental data to different thermodynamic equilibrium models without (n=1, left plot) and with (n=2, right plot) cooperativity gave an estimate of ~100-fold lower $K_D$ value (values in legend).

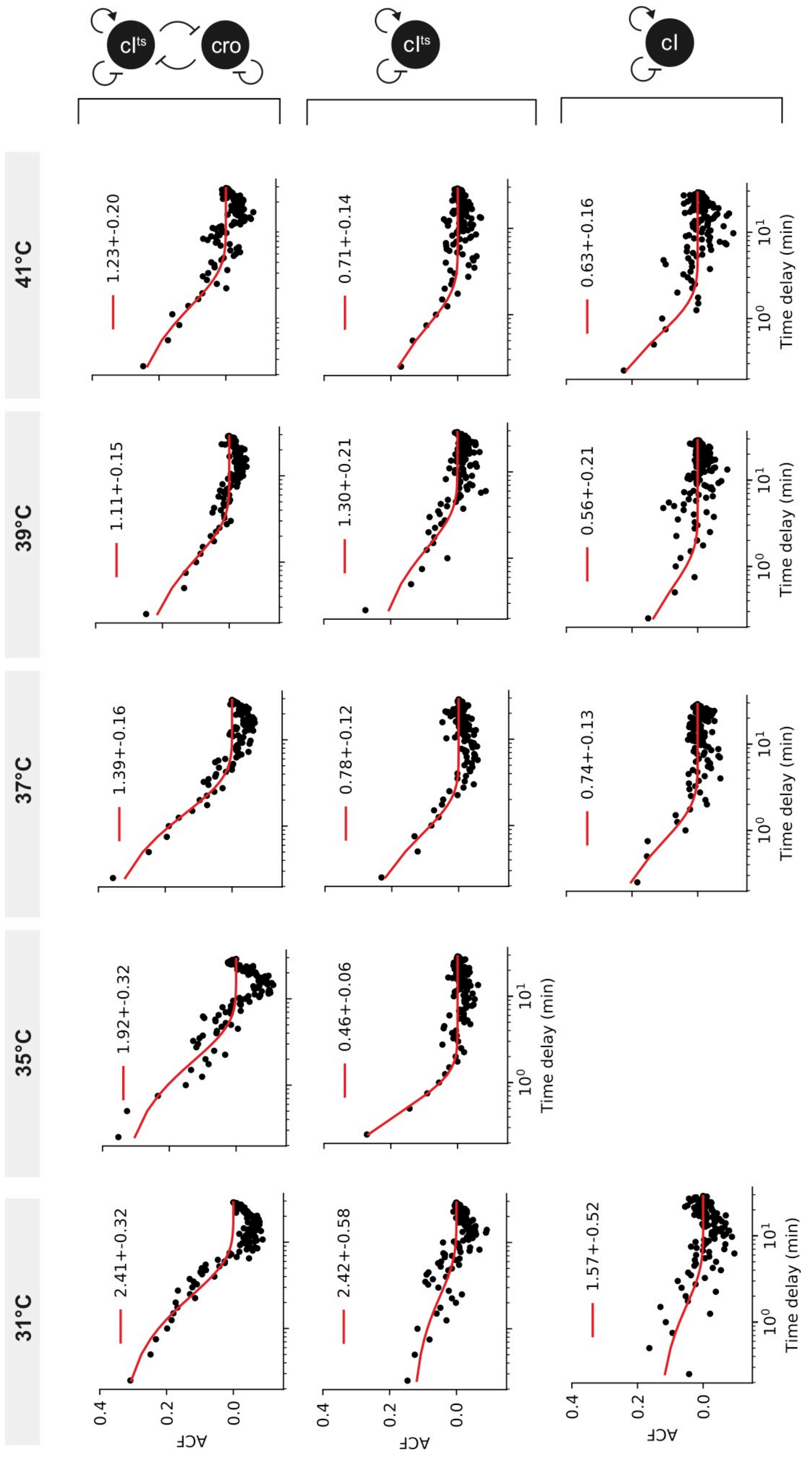

31°C
35°C
37°C
39°C
41°C
2.41+-0.32
1.92+-0.32
1.39+-0.16
1.11+-0.15
1.23+-0.20
2.42+-0.58
0.46+-0.06
0.78+-0.12
1.30+-0.21
0.71+-0.14
1.57+-0.52
0.74+-0.13
0.56+-0.21
0.63+-0.16
ACF
Time delay (min)
cI^ts
cro
cI

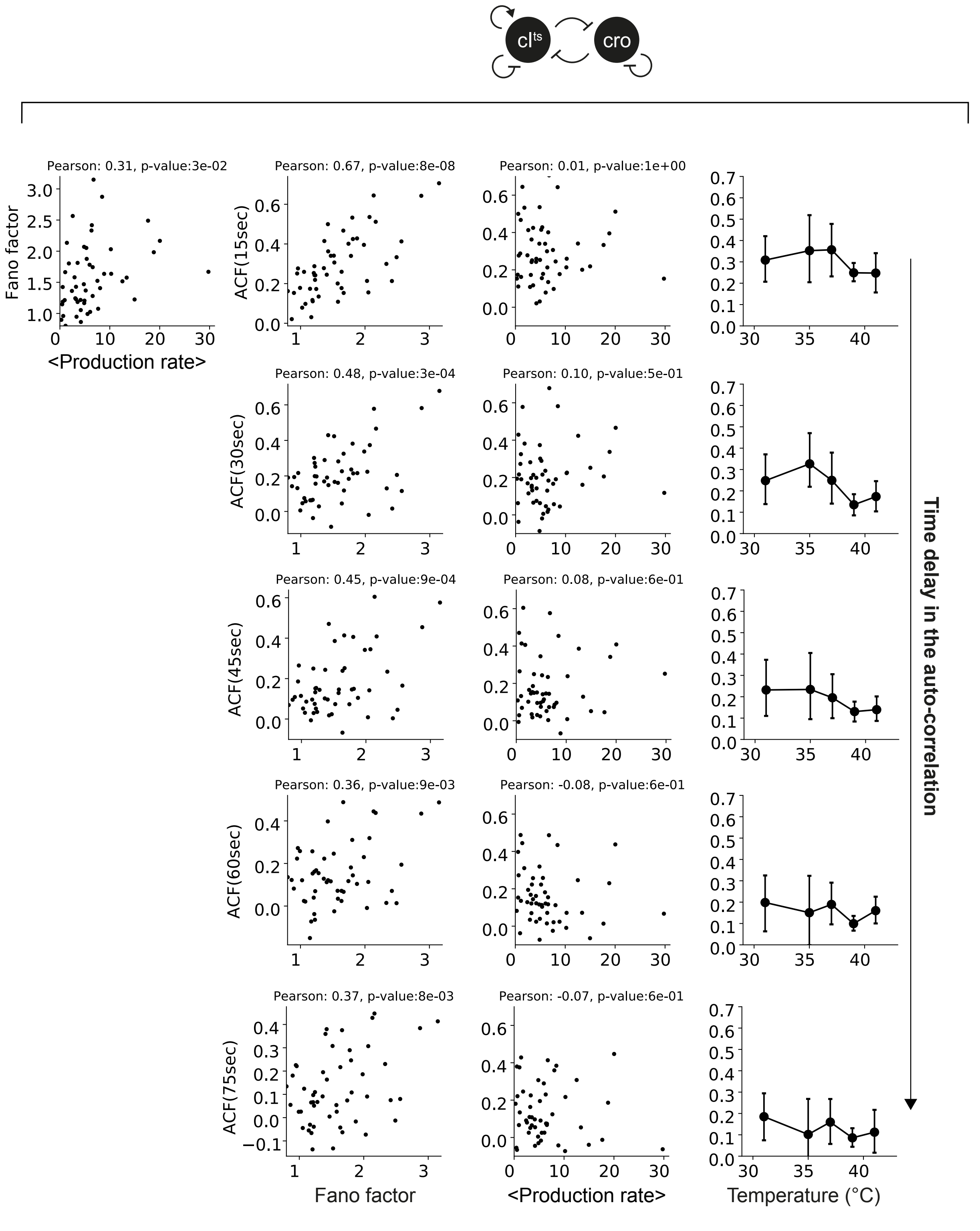
cl^ts
cro
Pearson: 0.31, p-value:3e-02
Fano factor
<Production rate>
Pearson: 0.67, p-value:8e-08
ACF(15sec)
Pearson: 0.01, p-value:1e+00
Pearson: 0.48, p-value:3e-04
ACF(30sec)
Pearson: 0.10, p-value:5e-01
Pearson: 0.45, p-value:9e-04
ACF(45sec)
Pearson: 0.08, p-value:6e-01
Pearson: 0.36, p-value:9e-03
ACF(60sec)
Pearson: -0.08, p-value:6e-01
Pearson: 0.37, p-value:8e-03
ACF(75sec)
Pearson: -0.07, p-value:6e-01
Fano factor
<Production rate>
Temperature (°C)
Time delay in the auto-correlation

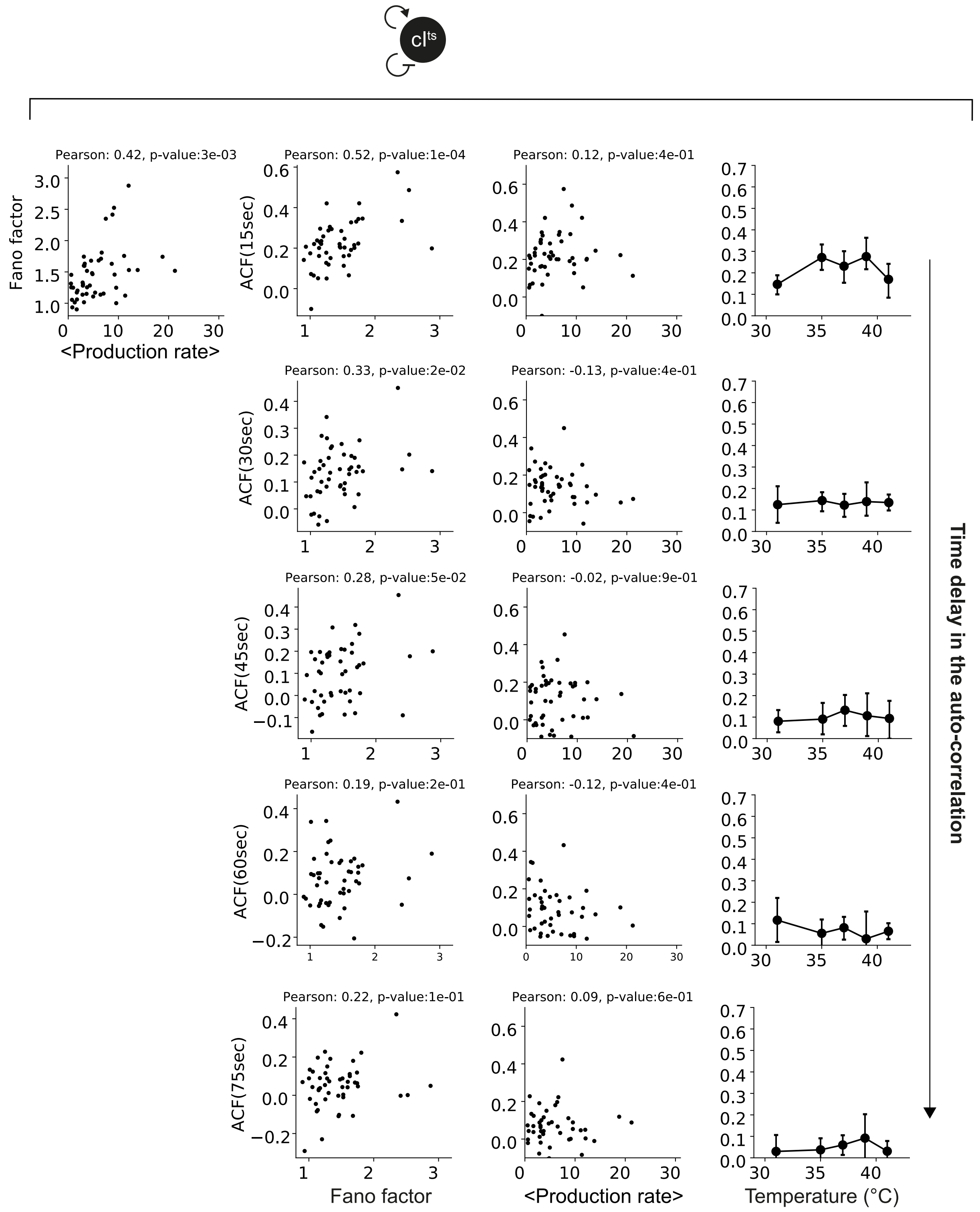


**Fig. S17. Ensemble-averaged autocorrelation function, correlations of time-averaged production rates, and Fano factor, and auto-correlation amplitudes at different time delays for the different GRNs.** First three columns show the ensemble-averaged ACFs from Fig. S12 at various temperatures and fits to a mono-exponential decay (red line) inspired by Yu *et al.* (23) for the three indicated GRNs (decay constant in minutes as show in figure legends). Center (bistable GRN) and bottom (monostable GRN) figure shows Fano factor against time-averaged production rates, ACF amplitude at the selected time delay against Fano factor, ACF amplitude at the selected time delay against time-averaged production rates, and ACF amplitude at the selected time delay against temperature (from left to right). Pearson correlation and p values are given in the title of each plot. The time delay is given in the y label in each GRN column (15, 30, 45, 60, 75 sec from top to bottom).

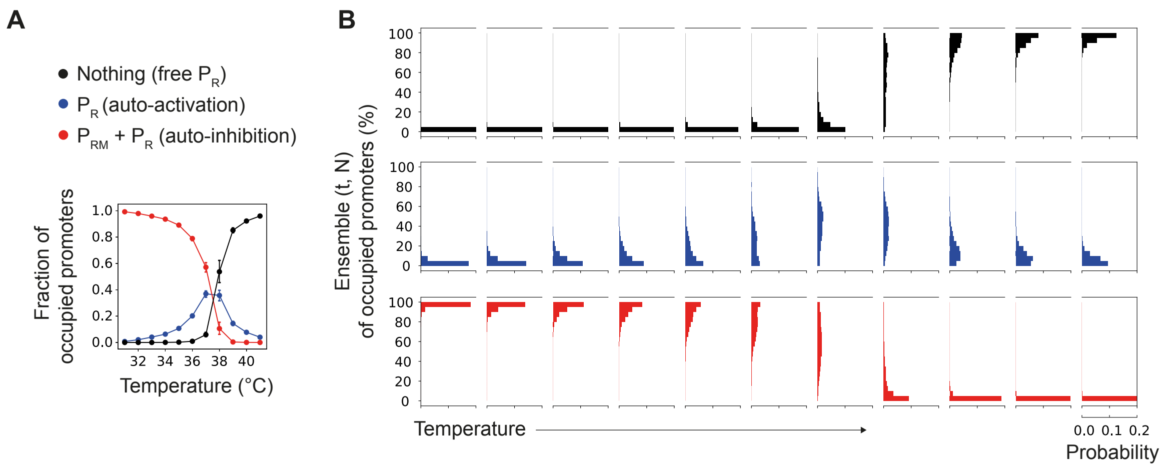


**Fig. S18. Simulated promoter occupancies.** (**A**) Averaged fraction of occupied promoters (time-averaged) by CI as obtained from simulation (as in Fig. 3D and Table S3 with $k_{ON}$=3e-15) for three different categories: No CI is bound to $P_R$ (black line), CI is bound to the $P_R$ promoter (blue line), and CI is bound to $P_R$ and $P_{RM}$ (red line). Errorbars were bootstrapped and indicate s.d. between compartments. (**B**) Simulated ensemble promoter occupancy as sampled over time t and compartment N at various temperatures (left to right: 1 °C steps from 31 °C to 41 °C). The occupancies are given for free $P_R$ (upper row), bound $P_R$ (center row), and bound $P_R$ and $P_{RM}$ (lower row). A transiently unoccupied (free) $P_R$ promoter would allow leaky production of Cro already at ~35 °C.

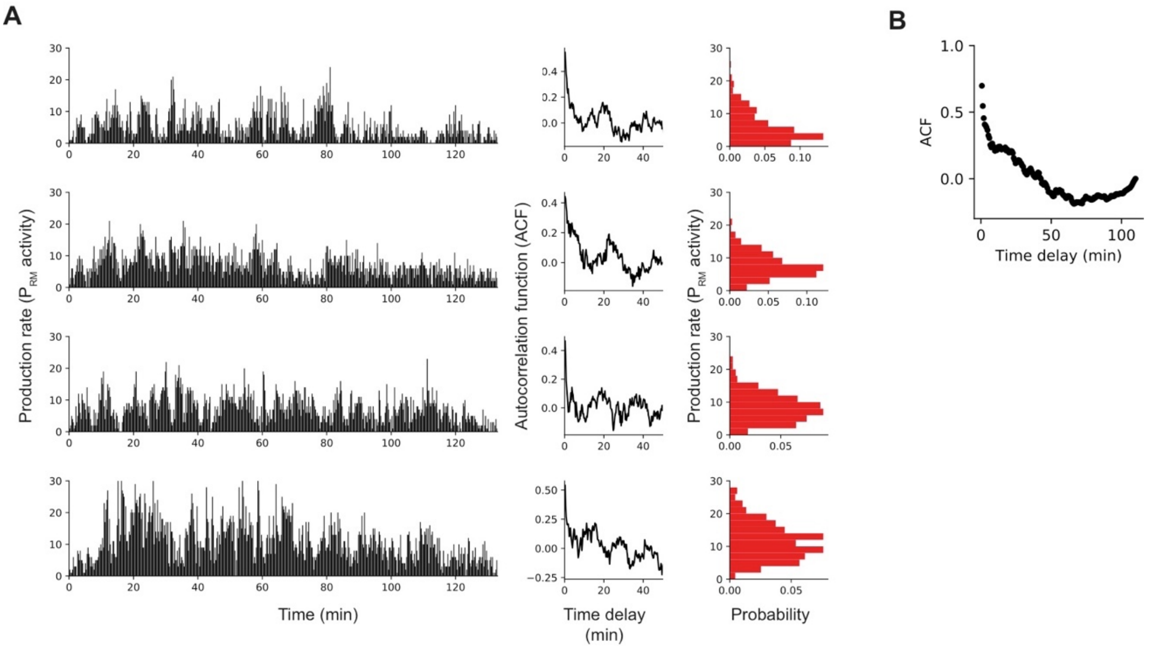


**Fig. S19. Long-term experiments of the bistable GRN at 37 °C.** (**A**) Individual production rates, individual ACFs, individual probability distributions of production rates (left to right) (**B**) Ensemble-averaged ACF of all compartments. The negative auto-correlation at 50-100 min may stem from slow degradation of DNA, also indicated by the slow decay of production rates in the individual production rates.

**Table S1. Primer sequences.**

| Name | Sequence | Mod | Usage |
|---|---|---|---|
| *P1* | GGAGATGGCGCCCAACAGTCGCAA TTCAGAGCGGCAGCAAGTG | Internal Modification | Circular DNA |
| *pP1* | GTGGATAACCGTATTACCGCCTTTG AGTG | 5'-Pi | Circular DNA |
| *P2* | GACTGTTGGGCGCCATCTCCGTGG ATAACCGTATTACCGCCTTTGAGTG | Internal Modification | Circular DNA |
| *pP2* | GCAATTCAGAGCGGCAGCAAGTG | 5'-Pi | Circular DNA |
| *P3* | CGCCGCAGAGTGGATGTACAGAAA AGCCCGCCTTTCGGCGGGCTTTGC TCGAGTTATCAGCCAAACGTCTCTT CAGGC | | Lift-off from lambda genome |
| *P4* | TATATCTCCTTCTTAAAGTTAAACAA AATTATTGCTAGCTTATGCTGTTGTT TTTTTGTTACTCGGGAAG | | Lift-off from lambda genome |

**Table S2. Ensemble averaged production rates for the various GRNs. The values are given in the following order: 68th *I median I* 32th.**

| Temperature (*℃)* | Bistable GRN | Monostable GRN | Wild-type monostable GRN |
|---|---|---|---|
| *41* | 6 / 4 / 3 | 5 / 4 / 3 | 6 / 5 / 4 |
| *39* | 10 / 7 / 4 | 10 / 7 / 5 | 6 / 4 / 3 |
| *37* | 9 / 6 / 4 | 12 / 9 / 7 | 5 / 3 / 2 |
| *35* | 8 / 5 / 3 | 5 / 3 / 2 | - |
| *31* | 1 / 0 / 0 | 1 / 1 / 0 | 1 / 0.5 / 0 |

**Table S3. Parameters for stochastic simulations.**

| *Parameters* | *Values* | *Units* | *Reference* |
|---|---|---|---|
| *Basal transcription rate* | 0.09 | 1/min | Estimated by the time-averaged production rate at 41 °C in the monostable GRN |
| *Auto-activated transcription rate* | 0.9 | 1/min | Estimated by the time-averaged production rate at 37 °C in the monostable GRN |
| $k_{on}$ *for* $O_R2$ *and* $O_R1$ | 3e-11 to 3e-15 | l/(#*min) | Lowest value taken from (28) |
| $k_{on}$ *for* $O_R3$ | 2e-11 to 2e-15 | l/(#*min) | Estimated from (28) and to integrate the fact that CI has a lower affinity to $O_R3$ than $O_R2$ and $O_R1$ (16) |
| $k_{off}$ | 0.25 | 1/min | ~1.5 1/min (28) |
| *Translation rate* | 1.3 | 1/min | ~3 proteins per mRNA was estimated and supported by (10) with 6 proteins per mRNA considering lower expression activity in cell-free extract |
| *mRNA degradation rate* | 0.5 | 1/min | (38) |
| *Amount of DNA* | 20 | # | This work |
| *Protein life-time and deactivation rate* | 0.04*exp(0.55*(T-32)) | 1/min | (19); We assumed a longer protein life-time (25 against ~10 min) due to hindered diffusion from DNA binding. |
| *Compartment volume* | 3.8e-12 | l | This work |
